# High Dose Gammaherpesvirus Infection in Macrophages Results in Inflammatory Multimodal Cell Death

**DOI:** 10.64898/2026.09.16.752237

**Authors:** Gabrielle Vragel, Gracyn Nelson-Reid, Rachael E. Kostelecky, Elizabeth A. Spear, Moses Lee, Derek W. Abbott, Linda F. van Dyk, Eric T. Clambey

**Author notes:** Co-Corresponding Senior Authors.

## Abstract

Human gammaherpesviruses (yHVs) including Epstein-Barr Virus and Kaposi’s sarcoma associated herpesvirus are linked to cancer and inflammatory disease development. The outcome of yHV infection is highly regulated by cell type, with macrophages identified as an important early infection target that can also serve as a latency reservoir. Here, we used the mouse yHV model, murine gammaherpesvirus (MHV68), to identify a dose-dependent outcome of MHV68 infection of macrophages, with high dose infection resulting in extensive morphological changes, viral gene expression, and a pronounced inflammatory cell death not observed under conditions of lytic replication occurring at lower infectious dose. Virion envelope and tegument components were not sufficient for this dose-dependent cell death, which required intact viral DNA and virus transcription, yet occurred independently of viral DNA replication and late gene transcription. Necrostatin-1 (Nec-1) treatment, an anti-inflammatory therapeutic, limited cell death and reduced cell morphology alterations, with no impact on virus replication, uncoupling cell death from lytic replication. Activation of autophagy profoundly limited high dose infection outcomes, with increased cell survival and reduced viral gene expression. Despite the improved cell survival observed in cells treated with Nec-1, biochemical analysis of infected cultures identified robust apoptosis induction with subsequent activation of gasdermin E, suggesting a multi-modal cell death mechanism culminating in pyroptosis. Our findings suggest that high dose macrophage infection byMHV68 is a major driver of inflammatory cell death to potentially shape downstream inflammation and immune responses.

**IMPORTANCE:** The human gammaherpesviruses (yHVs) are globally prevalent viruses which can cause many cancers and chronic inflammatory diseases. Despite their pervasiveness, little is known about the nature of yHV infection in macrophages, a relevant host cell type, that is highly responsive to the environment and plays major roles in immune modulation. During infection, macrophages are likely to be exposed to different amounts of virus, based on their proximity to sites of ongoing virus replication or reactivation. By studying how macrophages respond to different amounts of virus, we find that the amount of virus macrophages are exposed to determines the likelihood of survival versus an inflammatory death mediated by integrated programmed cell death mechanisms. This inflammatory death can be inhibited by Necrostatin-1, a promising treatment for multiple inflammatory diseases. Overall, our work highlights the impact of virus dose on promoting oncogenic conditions and introduces a potential treatment option to limit this inflammation.

## INTRODUCTION

The gammaherpesvirus (yHVs) are large oncogenic DNA viruses that include Epstein-Barr virus (EBV) and Kaposi’s sarcoma-associated virus (KSHV), and infect their hosts for life. The yHVs have a high prevalence in the human population, with EBV infecting over 90% of the global population, resulting in over 300,000 new cases of cancer in 2020 alone [1, 2]. Murine gammaherpesvirus 68 (MHV68 or yHV68; ICTV nomenclature, *Murid herpesvirus 4*, MuHV-4) is a small animal model of yHVs, with genetic and biologic similarity to the human yHVs, which affords detailed analysis of multiple stages of infection in vitro and in vivo [3, 4]. The outcome of yHV infection is biphasic and highly dependent on cell type. Lytic replication, characterized by full viral gene transcription and virus replication, is common in epithelial cells and fibroblasts. Latent infection, characterized by little to no viral gene expression and no production of infectious virions, occurs predominantly in B cells, a long-term virus reservoir [5]. Myeloid cells, including macrophages, monocytes and dendritic cells, have critical functions during yHV infection: they promote the early antiviral response yet are also targeted for infection and subject to viral manipulation by EBV, KSHV, and MHV68 [6–23]. In MHV68 and KSHV, myeloid cells also facilitate B cell infection [9, 21, 24].

Beyond host target cell type, yHV infection outcome is regulated by numerous virus and host derived factors. We recently reported that macrophages infected with an intermediate infection dose of MHV68 (1 plaque forming unit (PFU) per cell) were efficiently infected, with very limited lytic replication [20]. In contrast, infection with a high dose (10 PFU/cell) of MHV68 was able to overcome this host restriction, to initiate lytic gene expression and virus replication [20]. Infectious dose is well-known to impact the outcome of herpesvirus infection, including examples where viral mutant defects can be overcome with increased infection dose (i.e. a higher multiplicity of infection) [25–29].

Host cell survival during yHV infection is a complex process influenced by virus factors, whether infection is lytic or latent, and the host cellular milieu. Seminal early studies focused on virally-encoded homologs (e.g. viral Bcl2 and FLIP) that promoted cell survival [30–33], often studying these processes outside the context of infection. As knowledge of cell death pathways has expanded to encompass non-inflammatory (i.e. apoptosis) and inflammatory (e.g. pyroptosis, necroptosis, ferroptosis) mechanisms of death, and these processes are increasingly studied in the context of bona fide infection (e.g. [34–39]) there is significant interest in how yHV infection intersects with these pathways to achieve lifelong infection and to promote cancer and chronic inflammation, especially in immunosuppressed individuals.

Here we show that high dose MHV68 infection of macrophages leads to a dramatic decrease in cell viability of both cell lines and primary cells. We find that high dose infection-associated cell death is a distinct outcome, separable from other states of productive lytic replication in macrophages, requiring viral transcription but occurring independently of viral DNA replication and late gene transcription. Notably, high dose infection elicits an inflammatory cell death, potentially triggering apoptosis, which in turn leads to gasdermin E-driven secondary pyroptosis. High dose infection further results in pronounced cell morphology changes and the generation of large, virus-infected giant cells that are disrupted by the RIPK1 and IDO inhibitor necrostatin-1. These results reveal that MHV68 infection can induce profound, dose-dependent alterations to the outcome of macrophage infection and cell fate.

## RESULTS

### High dose (MOI=10) infection in J774 and peritoneal macrophages results in pronounced cell death

We previously found that high dose (MOI=10) MHV68 infection in the mouse J774 macrophage cell line resulted in productive replication accompanied by a high frequency of cell death, an outcome not observed with intermediate dose infection (MOI=1) [20]. To further investigate this dichotomy, we infected J774 cells with intermediate (MOI=1) or high dose (MOI=10) MHV68.H.GFP virus, a recombinant virus that robustly expresses GFP during lytic replication [40]. While cultures infected with intermediate dose had limited cytopathic effect (CPE) and few GFP+ cells by 48 hpi, high dose infection was characterized by cell swelling, bright GFP puncta, vacuolization, and multiple nuclei, with morphologic similarities to giant cells or ballooning cells seen in secondary necrosis [41, 42] (**Fig 1A, Fig S1A**) with a high frequency of GFP expression and a low frequency of cell viability **(Fig 1B-D, Fig S1B**). Cell death occurred in WT MHV68, and in wild-type reporter virus-infected cells (MHV68.H.GFP, MHV68.LANA::βlac), in a dose-dependent manner, with decreased cell viability observed at MOI=5 and maximal death observed following high dose (MOI=10) infection (**Fig S1C**). As peritoneal macrophages are a major in vivo target of primary infection, and a latency reservoir [10, 43], we tested infection outcomes in primary peritoneal macrophages. At 48 hpi, peritoneal cell viability remained high in mock and intermediate dose infection cultures, in contrast to high dose infection cultures which demonstrated pronounced cell death (**Fig 1E**). These data emphasize that cell death resulting from high dose infection occurs in both a macrophage cell line and primary macrophages infected ex vivo.

**Figure 1:**
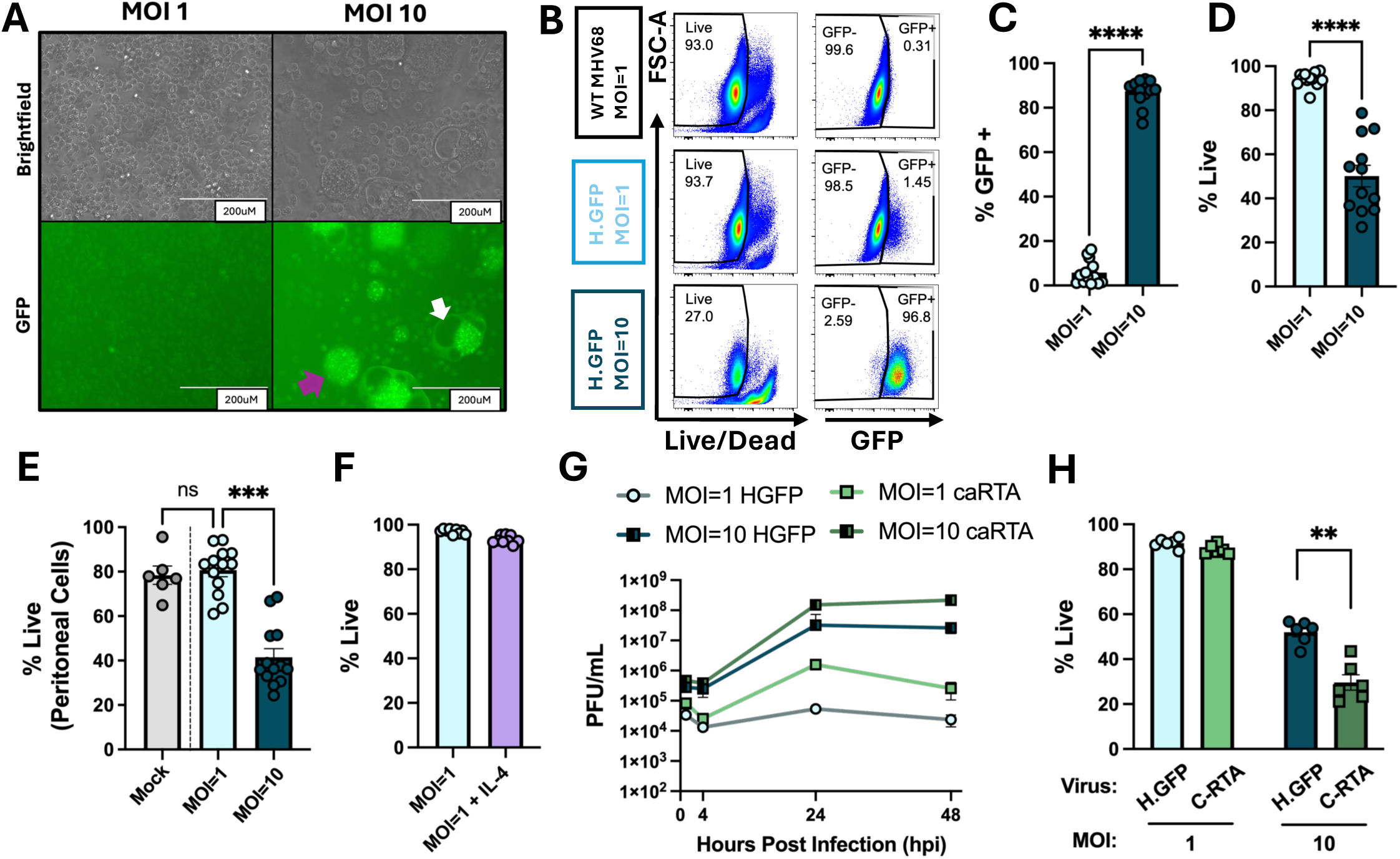
High dose (MOI=10) infection in J774 and peritoneal macrophages results in pronounced cell death. Analysis of virus infection outcome in MHV68 infected **(A-D, F-H)** J774 or **(E)** peritoneal macrophages as a function of different treatment conditions. **(A)** 20X Brightfield or corresponding GFP images of J774 macrophages infected with MHV68.H.GFP at MOI=1 or 10, harvested 48 hpi. White scale bar indicates 200 microns. Purple arrow indicates large, GFP+ cell. White arrow indicates vacuolization. **(B)** Representative flow cytometry plots, assessing the frequency of viable cells among single cells (i.e. Live, left column) and GFP+ cells among single, viable cells (right column) of J774 macrophages infected with MOI=1 or 10 of WT MHV68 or H.GFP virus, harvested at 48hpi. **(C-D)** Frequency of **(C)** GFP+ cells or (**D)** live cells among single cells, as in panel **B**, with full gating strategy in **Fig S1B**. **(E)** Frequency of live cells, defined by flow cytometry, in peritoneal cells harvested from C57BL/6Jmice and infected ex vivo with H.GFP virus at MOI=1 or 10, harvested at 48 hpi. **(F)** Frequency of live cells, defined by flow cytometry, of J774 macrophages infected with WT.LANA::βlac with or without 10ng/mL IL-4 pre-treatment, MOI=1, harvested at 48 hpi. **(G)** Virus replication defined by plaque assay, comparing J774 cells infected with MHV68.H.GFP or C-RTA with MOI=1 or 10, harvested at the indicated timepoint. **(H)** Frequency of live J774 macrophages infected with H.GFP or C-RTA virus at MOI=1 or 10. Flow cytometry-defined values **(B-F, H)** quantified the frequency of live cells among single cells (i.e. % live) and the frequency of GFP+ cells among single, viable cells (i.e. % GFP+), with full gating strategy in **Fig S1B**. Data show mean ± SEM, with individual symbols showing independent biological samples from 2-3 independent experiments with 2-3 biological replicates per experiment. Plaque assay measurements (**G**) were done in technical triplicates, with mean value shown for each biological replicate. Statistical analysis done using Mann-Whitney test (**C, D, H**) or one-way ANOVA (**E**, Kruskal-Wallis test subject to Dunn’s multiple comparisons test) with statistically significant differences indicated, **p<0.01, ***p<0.001, ****p<0.0001, ns (not significant).

Lytic herpesvirus infection in permissive cells is characterized by extensive CPE and cell death, often attributed to cell lysis [5]. As high dose infection in macrophages results in increased virus replication, we investigated whether other conditions associated with virus replication in macrophages were also characterized by high frequencies of cell death. First, we analyzed the impact of interleukin-4 (IL-4), a cytokine that promotes virus replication in macrophages [20, 44, 45]. Despite the robust ability of IL-4 to promote virus replication in vitro [20], J774 cells had no difference in viability comparing cultures infected with intermediate dose MHV68 in the presence or absence of IL-4 (**Fig 1F**). Next, we tested the consequence of infection with MHV68.C-RTA, a virus that overexpresses the viral immediate early transactivator, RTA, robustly initiating lytic replication in vitro and in vivo [46]. While MHV68.H.GFP-infected cultures showed no discernable infectious virus production by plaque assay, C.RTA-infected cultures showed abundant virus production at intermediate dose, with no decrease in cell viability (**Fig 1G-H**). Cultures infected with high dose C.RTA had increased virus production with lower cell viability than H.GFP-infected cultures but less than high dose infection (**Fig 1G-H**). These data indicate that high dose MHV68 infection in macrophages results in a dramatic cell death that is not observed under other conditions of virus replication in J774 cells.

### Cell death resulting from high dose infection of macrophages requires intact viral DNA and occurs independently of viral late gene transcription and viral DNA replication

We next sought to define how distinct infection stages contributed to high dose-associated cell death. To investigate whether cell death required intact, infectious virus particles, we tested the infection outcome with MHV68 inactivated with centanamycin (CM), a dsDNA-specific alkylator that has previously been used to inactivate multiple herpesviruses (HSV, HCMV, MCMV) [47]. Importantly, CM-treatment specifically inactivates viral DNA without altering viral envelope, tegument, and capsid proteins, allowing us to test the contribution of incoming virus proteins on high dose infection-associated death. Viral stocks were treated with either vehicle or 100 µM CM for 2 hours, after which virus was centrifuged, washed and tested for infectivity by multiple measures (**Fig 2A**). As expected, vehicle-treated samples had near WT levels in virus titer (**Fig S2A**). In contrast, CM-treated virus stocks showed negligible infection following infection with WT MHV68.LANA::βlac (**Fig 2B-C**), a sensitive reporter virus that measures the frequency of cells expressing the MHV68 LANA protein (**Fig S2B**), which is expressed during lytic and latent infection [48]. CM-treated virus stock was inactivated by multiple metrics, including no detectable infectious virus when tested on mouse embryonic fibroblast monolayers (**Fig S2C**), and negligible frequencies of GFP expression in 3T12 cells infected with either MHV68.ORF59.GFP (MOI=1) or MHV68.H.GFP (MOI=9) (**Fig 2D-E**). Virus inactivation was ineffective at lower CM concentrations or following CM treatment at 4° C (**Fig S2D**). These data demonstrate that CM can effectively inactivate MHV68, preventing transcription and replication of viral DNA in infected cells.

**Figure 2:**
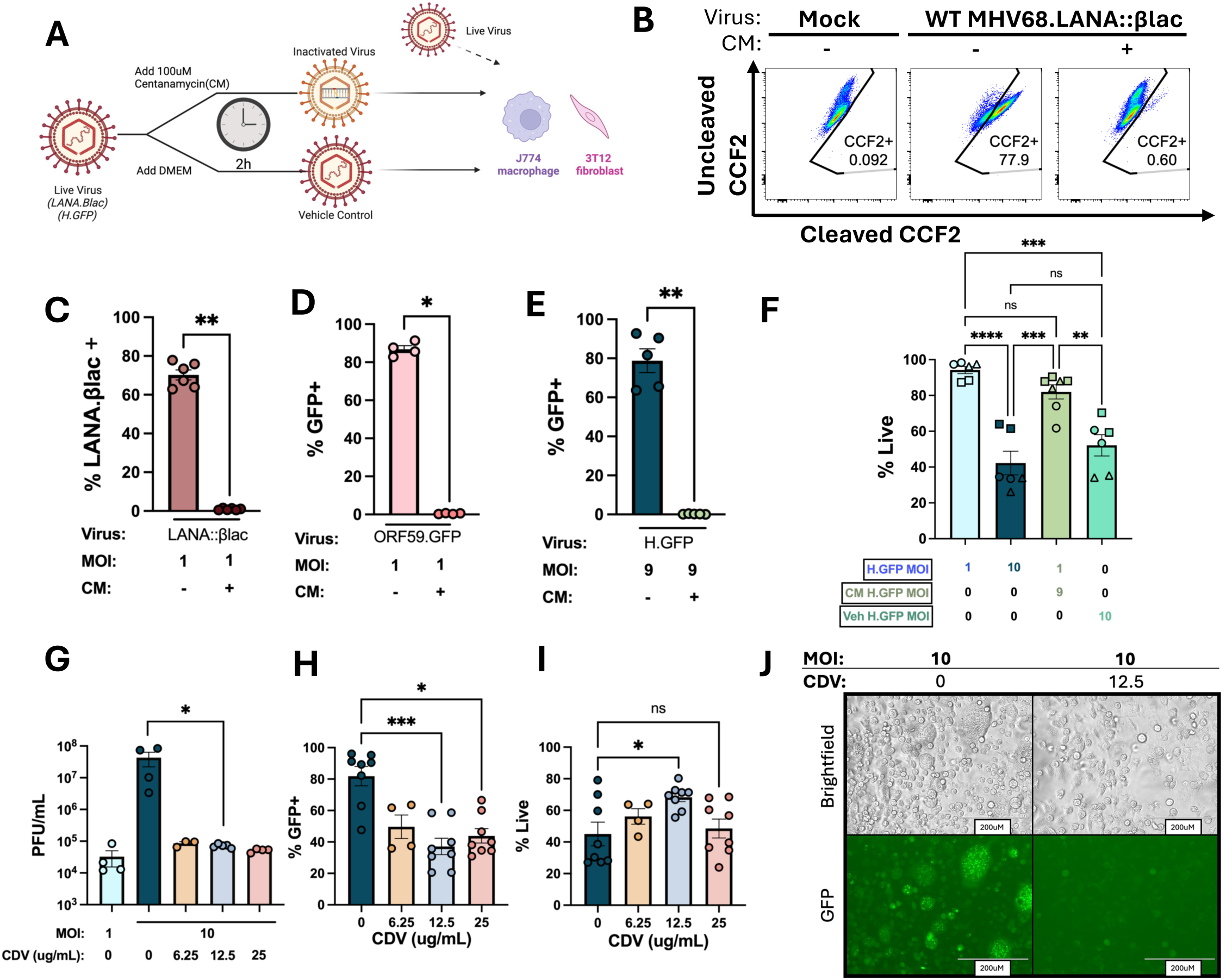
Cell death resulting from high dose infection of macrophages requires intact viral DNA and occurs independently of viral late gene transcription and viral DNA replication. Analysis of outcomes of high MOI infection in J774 macrophages as a function of virus treatment. (A) Schematic of virus inactivation by centanamycin (CM). 100 µM CM or vehicle was added to live virus and incubated for 2h at room temperature. Virus was spun down and resuspended in original volume, titered using the vehicle control, with virus inactivation defined in 3T12 or J774 cells, comparing mock or WT.LANA::βlac infected cells. (B,C) Frequency of MHV68-infected, LANA.βlac+ cells defined by flow cytometry, detected by CCF2 substrate cleavage, analyzing single, viable cells at 48 hpi, comparing mock or WT.LANA::βlac infected 3T12 fibroblasts (MOI=1), illustrated by (B) representative flow cytometry plots with (C) corresponding quantitation. (D-E) Frequency of viral gene expression defined by flow cytometry, in (D) 3T12 cells infected with MHV68.ORF59.GFP (MOI=1) harvested at 48 hpi, and in (E) 3T12 cells infected with MHV68.H.GFP (MOI=9) harvested at 24 hpi. (F) Frequency of live J774 cells infected with H.GFP with or without CM treatment, harvested at 48 hpi, defined by flow cytometry, with shapes indicating paired experimental samples. (G) Virus replication in J774 macrophages infected with H.GFP with or without cidofovir (CDV) treatment at various doses. (H-I) J774 cells were treated with CDV 1h pre-infection at indicated dose, followed by infection with MOI=10 H.GFP. Frequency of (H) GFP+ or (I) live cells defined by flow cytometry at 48 hpi. (K) 20X brightfield or corresponding GFP images of CDV treated or untreated J774 macrophages at 48 hpi. White scale bar indicates 200 microns. % LANA.βlac+ events (B-C) defined by the frequency of CCF2 substrate cleavage among single, viable cells by flow cytometry using a 405 nm laser. CCF2 substrate cleavage defined by fluorescent emission (detected by 460-22 nm filter) relative to substrate loading (detected by 525-50 nm filter). Flow cytometry-defined values (D-F, H-I) quantified the frequency of live cells among single cells (i.e. % live) and the frequency of viral reporter positive cells among single, viable cells (i.e. % LANA.βlac+ or GFP+), with full gating strategy in Fig S2A. Data show mean ± SEM, with individual symbols showing independent biological samples from 2-4 independent experiments with 2-3 biological replicates per experiment. Statistical analysis was done using Mann-Whitney test (C-E, G), two-way ANOVA (F, ordinary, with Tukey’s multiple comparisons test), or one-way ANOVA (H, I). In panels G-I, statistical analysis only compared samples derived from at least three independent experiments (G, MOI=10 and 12.5 µg/mL CDV; H-I, MOI=10, 12.5 µg/mL, 25 µg/mL CDV). Statistically significant differences indicated, *p<0.05, **p<0.01, ***p<0.001, ****p<0.0001, ns (not significant).

The early stages of herpesvirus infection are characterized by viral glycoproteins binding to the host cell, followed by introduction of viral tegument proteins into the cytosol after viral fusion with the cell membrane [49]. In the herpesvirus field, there is also a body of literature identifying infection outcomes that can be complemented *in trans* by co-administration of inactivated virus, effects mediated by viral glycoprotein or tegument delivery by an inactivated virus particle [42, 50–52]. We therefore tested the outcome of intermediate dose infection (MOI=1) in which the virus inoculum was supplemented with high dose CM-inactivated MHV68 (MOI=9). Though high dose infection (MOI=10) with MHV68.H.GFP or vehicle-treated H.GFP virus resulted in pronounced cell death relative to intermediate dose infection (MOI=1), intermediate dose infection supplemented with an MOI=9 of CM-treated MHV68 did not result in the same magnitude of cell death observed with high dose infection (**Fig 2F**). In parallel, intermediate dose infection supplemented with an MOI=9 of CM-treated MHV68 did not significantly increase the frequency of GFP expressing cells above that observed with intermediate dose infection alone, dramatically lower than the frequency of GFP expressing cells observed with high dose infection (**Fig S2E**). As CM induces DNA alkylation, without degrading viral proteins (measured by the ability to induce antibodies against CM-treated virus particles [47]), these data suggest that delivery of viral envelope, tegument and capsid proteins are not sufficient to enhance cell death following intermediate dose infection. These data suggest that intact viral DNA, potentially allowing virus transcription, is required for high dose infection-associated cell death.

We next assessed how viral DNA replication and late viral gene transcription contribute to high dose infection-associated death, using cidofovir (CDV), an inhibitor of the viral DNA polymerase and late herpesvirus gene expression [53]. CDV treatment significantly reduced infectious virus production following high dose infection of J774 cells **(Fig 2G)** and reduced the frequency of H.GFP expressing cells at all concentrations of CDV tested (**Fig 2H**). In contrast, cell viability only modestly increased at 12.5 µg/mL CDV **(Fig 2I)**. CDV treatment also markedly reduced CPE, seemingly eliminating the development of large, highly GFP+ swollen cells (**Fig 2J).** The minor protection from cell death provided by CDV suggests that high dose infection-associated cell death occurs independently of viral DNA replication and late gene transcription. In sum, these data suggest that high dose MHV68 infection of macrophages results in cell death that requires intact viral DNA and some degree of virus transcription, independent of viral DNA replication and late gene transcription.

### High dose MHV68 infection of macrophages results in inflammatory cell death inhibited by Necrostatin-1

Programmed cell death can be broadly divided into inflammatory or non-inflammatory cell death. Based on the extensive morphological changes observed after high dose infection of J774 macrophages, we investigated whether cells underwent an inflammatory cell death, as measured by lactate dehydrogenase (LDH) release [54–56]. Studies focused on necroptosis and pyroptosis, two inflammatory programmed cell death pathways implicated in other herpesvirus infections [37, 57–60]. First, we tested the susceptibility of J774 cells to necroptosis and pyroptosis inducers, as well as inhibitors of these pathways, using Necrostatin-1 (Nec-1, a receptor interacting protein kinase 1 (RIPK1) inhibitor [61, 62]) and LDC7559 (LDC, a Gasdermin D (GSDMD) inhibitor [63]). J774 cells treated with known necroptosis activators (TNFα, the caspase inhibitor Z-VAD-FMK, and the Smac mimetic sm-164 [64]) resulted in increased cytotoxicity, measured by LDH release, an outcome blunted by Nec-1 treatment **(Fig 3A)**. In parallel, J774 cells treated with lipopolysaccharide (LPS, an inflammatory pathway stimulator) and nigericin (driving potassium efflux to assemble the inflammasome) resulted in increased cytotoxicity which was reduced by LDC7559 **(Fig 3A)**. These data show that uninfected J774 cells are susceptible to necroptosis and pyroptosis, both processes which can be inhibited by drug treatments.

**Figure 3:**
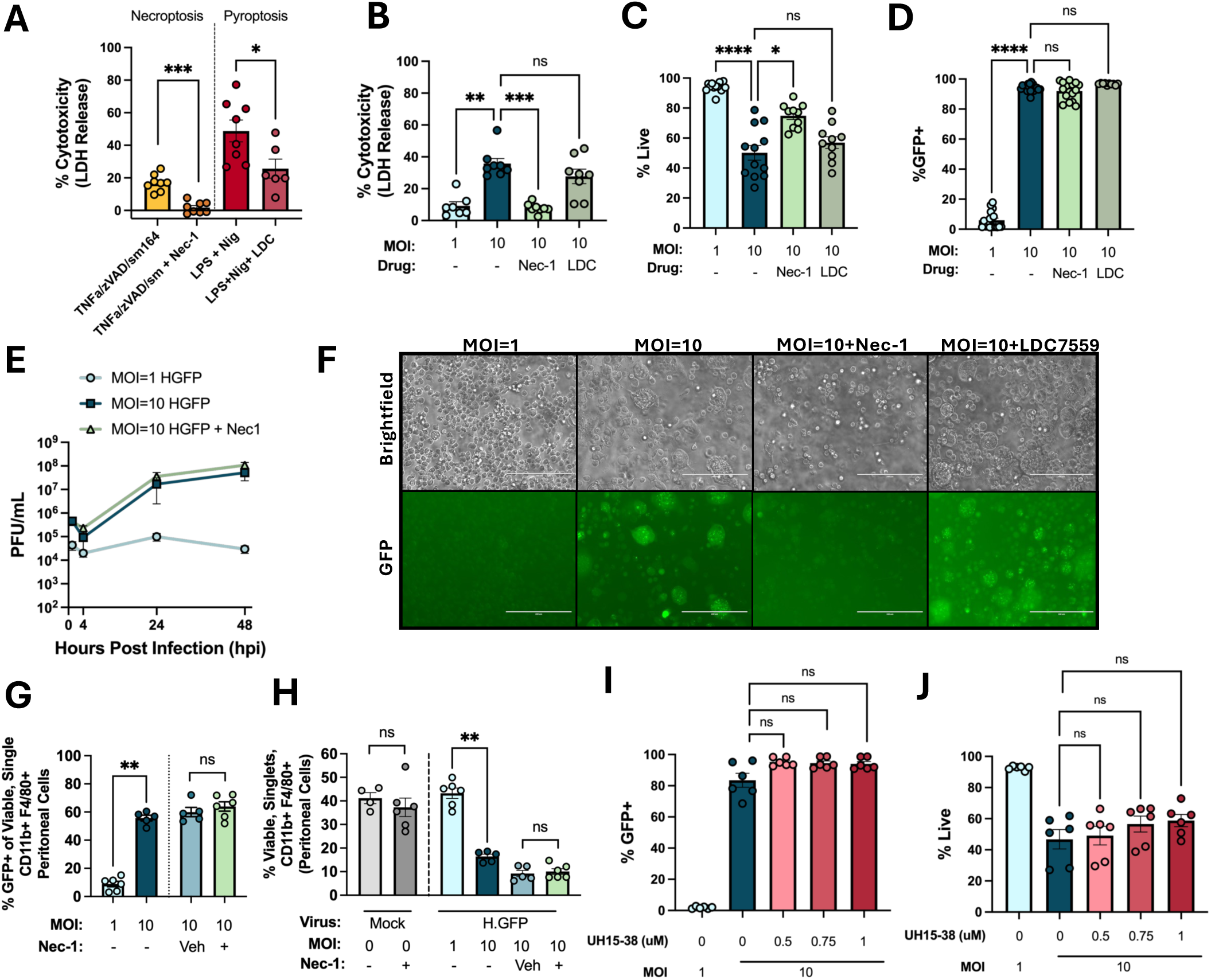
High dose MHV68 infection of macrophages results in inflammatory cell death inhibited by Necrostatin-1. Investigation of cell death mechanisms in MHV68-infected (B-F, I-J) J774 macrophages or (F-G) peritoneal cells. (A) LDH release from J774 macrophages as a measure of cytotoxicity when treated with necroptosis activators [TNFα (25 ng/mL), z-VAD-FMK (50 µM), and sm164 (5 µM)], with or without necroptosis inhibitor Nec-1 (30 µM) in 10% cDMEM (left) or pyroptosis activators [LPS (1 µg/mL) and Nigericin (Nig, 20 µM)], with or without pyroptosis inhibitor, LDC7559 (1 µM) in LCIS media. (B) LDH release from MHV68.H.GFP-infected J774 macrophages, at indicated MOI, treated with Nec-1 (30 µM) or LDC7559 (1 µM), harvested at 48 hpi. (C-D) Frequency of (C) live cells or (D) GFP expression in H.GFP-infected J774 macrophages with the indicated MOI, with or without treatment with Nec-1 (30 µM) or LDC7559 (1 µM) treatment, defined by flow cytometry at 48 hpi. (E) Virus titer by plaque assay, comparing virus replication as a function of MOI with or without Nec-1, harvested at indicated timepoints. (F-G) Frequency of (F) GFP expressing or (G) live H.GFP-infected peritoneal cells fromC57BL/6J mice, harvested and infected with H.GFP virus ex vivo at MOI=1 or 10 ± 30 µM Nec-1 treatment, harvested at 48 hpi. (H) 20X brightfield (top) or corresponding GFP images (bottom) of MHV68-infected J774 macrophages subjected to the treatment indicated above each column. White scale bar indicates 200 microns. (I-J) Frequency of (I) GFP expression or (J) live cells in H.GFP-infected J774 macrophages with the indicated MOI, with or without treatment with UH15-38, a RIPK3 inhibitor, 1h prior to infection with MHV68.H.GFP virus (MOI=1 or 10). (C-D, I-J) Data quantify the frequency of live cells among single cells (i.e. % live) and the frequency of GFP+ cells among single, viable cells or (F-G) the frequency of single, viable, CD11b+ F4/80+ peritoneal macrophages, with full gating strategy in Fig S3B. Data show mean ± SEM, with individual symbols showing independent biological samples from at least 3 independent experiments with 2-3 biological replicates per experiment. Plaque assay measurements (**E**) were done in technical triplicates, with mean value shown for each biological replicate. Statistical analysis was done using Mann-Whitney test (**A**, **G**, **H**) comparing the indicated pairs or one-way ANOVA (B-D, I-J, Kruskal-Wallis test subject to Dunn’s multiple comparisons test). In panels I-J, statistical analysis compared the effect of UH15-38 dose relative to MOI=10 without drug treatment. Statistically significant differences indicated, *p<0.05 **p<0.01, ***p<0.001, ****p<0.0001, ns (not significant). LDC, LDC7559; LPS, lipopolysaccharide; Nec-1, necrostatin-1; Nig, nigericin.

We next assessed the impact of intermediate and high dose MHV68 infection on cell death, following MHV68.H.GFP infection. By 48 hpi, high dose infection resulted in a significant increase in LDH release, indicative of inflammatory cell death, relative to intermediate dose infection (**Fig 3B**). High dose infection cytotoxicity was significantly reduced by Nec-1, but not LDC treatment (**Fig 3B**), with increased cell viability in Nec-1, but not LDC-treated cultures (**Fig 3C**). High dose infection resulted in high frequencies of GFP expressing cells that were not altered by Nec-1 or LDC treatment (**Fig 3D**); analysis of GFP fluorescence in Nec-1 treated cultures may be an over-estimate given the apparent autofluorescence in these cultures (**Fig S3A**). Nec-1 treatment did not affect the kinetics or magnitude of lytic replication following high dose infection (**Fig 3E**) but reduced large ballooning cells in high dose infection cultures with or without LDC (**Fig 3F**). These data indicate that in J774 cells, high dose infection results in an inflammatory cell death that is limited by Nec-1 treatment. While Nec-1 treatment limits cell death and reduces prominent cell morphology changes, it had no detectable impact on viral gene expression or the lytic replication in high dose infection conditions.

We previously found high dose infection induced cell death in primary peritoneal cells infected ex vivo (**Fig 1E**). We next analyzed the impact of Nec-1 on this infection outcome, using ex vivo infection of primary peritoneal cells. High dose infection increased the frequency of GFP expressing peritoneal macrophages relative to intermediate dose infections; neither vehicle or Nec-1 treatment altered the frequency of GFP expressing cells (**Fig 3G, Fig S3B**). Ex vivo cultured peritoneal macrophages retained high viability in mock- and intermediate dose infection conditions, with significantly reduced viability in high dose infection (**Fig 3H**). Cell viability was further reduced in both vehicle and Nec-1 treated cultures of high dose infection (**Fig 3H**).

These data indicate that while high dose infection resulted in increased cell death in J774 cells and peritoneal macrophages, the protective effects of Nec-1 were observed in J774 cells, not in ex vivo cultures of primary peritoneal macrophages.

Necroptosis involves multiple downstream mediators, including RIPK1, RIPK3 and MLKL [65]. Given that Nec-1, a RIPK1 inhibitor, was able to limit high dose-infected induced death, we next tested UH15-38 [66], a RIPK3 inhibitor. UH15-38 treatment had no effect on viability of mock infected J774 cells (**Fig S4A**). In contrast to the protective effects of Nec-1, UH15-38 treatment had no discernible effect on the frequency of GFP expressing cells or on cell viability following high dose infection of J774 cells (**Fig 3I-J**). In addition, we tested the contribution of TNFα to high dose infection-associated cell death; pretreatment of cultures with TNFα-blocking antibody had no effect on the frequency of GFP expressing cells or cell viability (**Fig S4B-C**). These data suggest either that this cell death pathway does not involve canonical necroptosis and/or that Nec-1 may be limiting cell death through RIPK1-independent effects (e.g. through its effects on IDO [67, 68]).

### High dose MHV68 infection of macrophages results in multimodal cell death with notable GSDME cleavage

We next investigated the effects of two viral genes previously implicated in regulating cell survival following MHV68 infection in endothelial cells, the MHV68 viral cyclin (vCyc) and viral Bcl2 (vBcl2) [34]. High dose infection with a vCyc-deficient MHV68 (vCyc.stop) resulted in equivalent cell death and LDH release to that seen with high dose WT infection (**Fig 4A-B**). In contrast, high dose infection with vBcl2-deficient MHV68 (vBcl2.stop [69]) that lacks the ability to inhibit apoptosis and autophagy, or with a vBcl2-mutant MHV68 (vBcl2.AAA [35]) that lacks the ability to inhibit autophagy, had reduced cell viability relative to high dose WT infection, with no difference in LDH release (**Fig 4A-B**). These data implicate MHV68 vBcl2 as one viral gene that promotes cell survival following high dose infection of macrophages.

**Figure 4:**
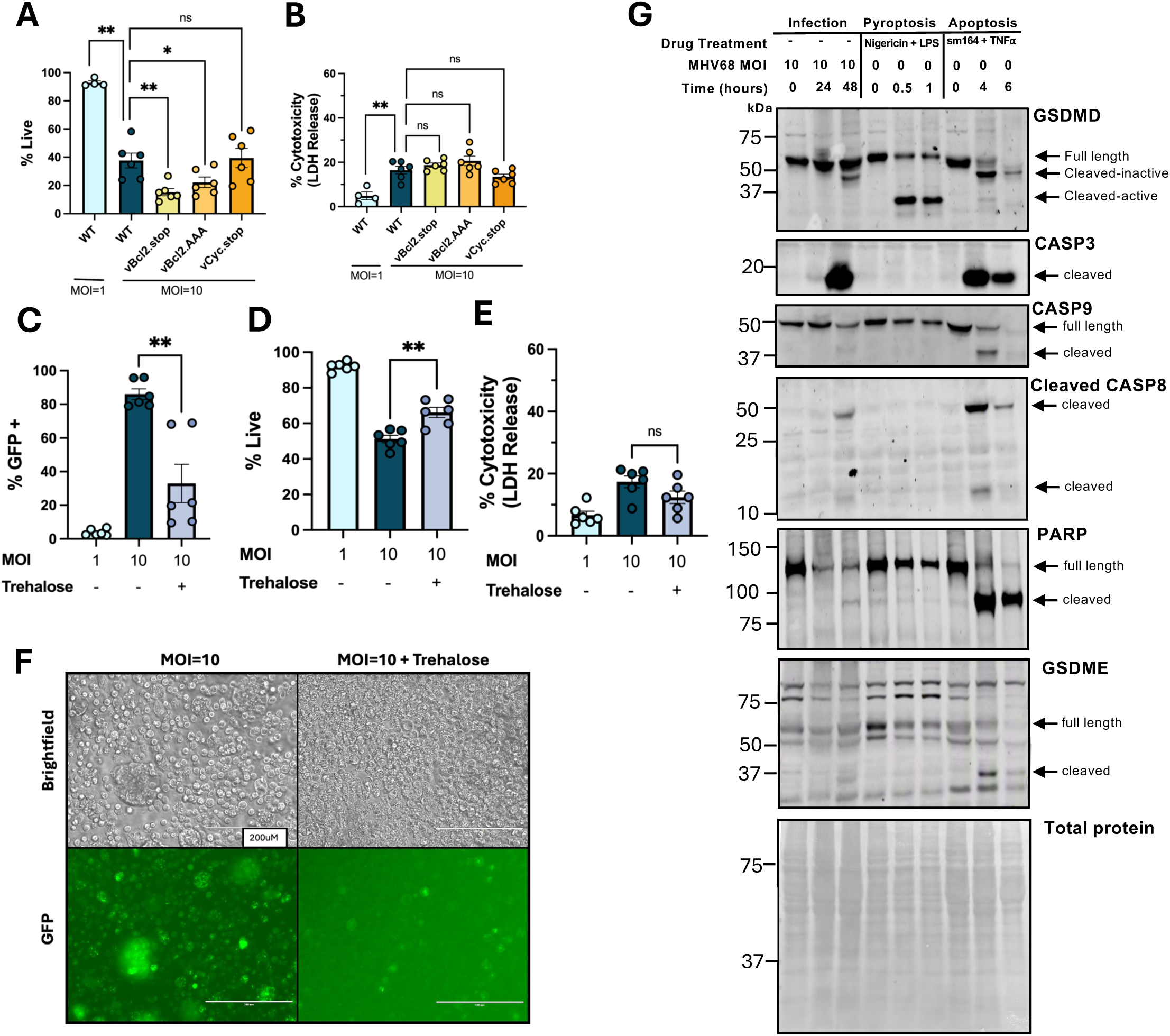
High dose MHV68 infection of macrophages results in multimodal cell death. Analysis of virus infection outcomes in J774 macrophages as a function of viral genes, autophagy and biochemical analysis of cell death pathways. (A-B) J774 macrophages were infected with indicated MOI of WT MHV68, vBcl2.stop, vBcl2.AAA or vCyc.stop viruses, harvested at 48 hpi. (C-E) J774 macrophages were treated with 100mM Trehalose during infection with MHV68.H.GFP (MOI=1 or 10), harvested 48 hpi. Frequency of (C) GFP expressing or (E) live cells measured by flow cytometry, with LDH release to measure cytotoxicity. (F) 20X brightfield or corresponding GFP images of MHV68.H.GFP-infected J774 macrophages untreated (left) or treated (right column) with trehalose, analyzed at 48hpi. White scale bar indicates 200 microns. (G) Immunoblot analysis of cell lysates from infected J774 cells (left) or J774 cells stimulated with pyroptosis inducers (nigericin with LPS, middle) or apoptosis inducers (sm164 with TNFα, right), probed with the indicated antibody; arrows on right side of each blot indicate expected sizes of cleaved protein products. Total protein loading measured by ponceau S staining is provided below; blot images shown are representative of three independent experiments with consistent results. Flow cytometry-defined values (A, C-D) quantified the frequency of live cells among single cells (i.e. % live) and the frequency of GFP+ cells among single, viable cells (i.e. % GFP+), with full gating strategy in Fig S1B. Data show mean ± SEM, with individual symbols showing independent biological samples from 3 independent experiments with 2 biological replicates per experiment. Statistical analysis was done using Mann-Whitney test (A-E) comparing either mutant virus (A-B) or the effect of trehalose (C-E) compared to MOI=10 infection conditions. Statistically significant differences indicated, *p<0.05 **p<0.01, ns (not significant).

A previous report demonstrated that the vBcl2.AAA mutant lacks the ability to limit autophagy [35]. We next tested the consequence of autophagy induction on high dose infection, treating cells with the autophagy-inducing compound trehalose.

Treatment with trehalose significantly decreased the frequency of GFP expressing cells in high dose infection conditions, increasing cell viability, visibly decreasing cell morphology changes, with no change in LDH release (**Fig 4C-F**). These results suggest that the intentional induction of autophagy in J774 cells alters the outcome of high dose infection, limiting fluorescence of a viral GFP transgene, increasing cell viability and attenuating cell morphology-associated changes.

We next sought to investigate biochemical features of cell death pathways in high dose infection, investigating indicators of pyroptosis, necroptosis, and apoptosis. While Gasdermin D (GSDMD), a pyroptotic effector mechanism [70], is expressed in J774 cells, and known pyroptosis-inducing stimuli resulted in cleavage to the active 30kDa form that results from Caspase 1; high dose infection did not result in active cleaved GSDMD, but instead showed only the 43kDa product that results from Caspase 3 cleavage and produces an inactive, truncated form of GSDMD [71] associated with apoptotic death (**Fig 4G**). Further, infected cell lysates had no evidence of MLKL phosphorylation, a terminal effector of necroptosis (**Fig S4E**), in agreement with resistance to RIPK3 inhibition, despite Nec-1 susceptibility (**Fig 3**). In contrast, high dose infection cultures were characterized by multiple measures of apoptosis induction, including robust induction of cleaved, active caspase 3, with lesser fractions of cleaved caspase 8, caspase 9 and PARP (poly-ADP-ribose polymerase) relative to cells undergoing apoptosis (**Fig 4G, Fig S4D**).

While high dose infection was clearly associated with the induction of apoptotic effectors, including caspase-3 activation, this finding alone did not explain the pronounced LDH release observed in high dose infected cells (**Fig 3B**). Since LDC7559 inhibits GSDMD, and we found no evidence for GSDMD activation (**Fig 4G**), we next examined the levels of gasdermin E (GSDME), another pyroptosis effector mechanism that can be activated by caspase 3, converting apoptotic to pyroptotic cell death [70, 72–74]. Activated GSDME protein products were detected in J774 cells induced to undergo apoptosis by 4 hours post-induction (**Fig 4G**), consistent with known caspase 3 cleavage. Additionally, high dose-infection cell lysates at 48hpi showed evidence of similar GSDME cleavage and activation (**Fig 4G**). MHV68 infection failed to robustly induce cell death in a series of immortalized bone marrow-derived macrophage cell lines (**Fig S4F**) reported to not express GSDME [73]. Taken together, these data suggest that high dose infection-associated cell death in J774 cells is multi-modal, inducing apoptotic machinery coupled with secondary necrosis, not through necroptosis or GSDMD, but potentially via Caspase 3 cleavage and subsequent GSDME-associated pyroptosis. MHV68 high dose, multimodal cell death can be limited by the vBcl2 and through the intentional induction of autophagy.

### High dose infection of J774 macrophages reveals dramatic cell morphology changes over time

To investigate the kinetics and nature of high dose infection-induced outcomes, we analyzed cell morphology, GFP expression and cell death over time using the Incucyte platform (supported by **Video S1-7)**. Mock and intermediate dose infection cultures were both characterized by a low frequency of dying cells (red cells, **Fig 5A**) with intermediate dose infection characterized by a low frequency of GFP expressing cells (green cells, second column from left, **Fig 5A**; **Video S1-2**). In striking contrast, high dose infection alone revealed an increased frequency of GFP expressing cells (12 hpi), which progressed to extensive GFP expression by multiple cells (by 24 hpi), followed by the coalescence of large, ballooning GFP expressing cells (by 36 hpi), after which a fraction of these cells progressed to cell lysis (by 48 hpi) (**Fig 5A** third column from left, **Fig 5B, Video S3-5**). In some cases, GFP+ ballooning cells appeared to fuse with other giant cells (e.g. **Fig 5A** blue outlined white arrow, 36 hpi). In other cases, GFP+ ballooning cells showed vacuolization (e.g. **Fig 5A** black outlined white arrow, 48 hpi, **Fig 5C**) or interacted with dead or dying cells (e.g. **Fig 5A** purple outlined white arrows, 36 hpi). High dose infection cultures had evidence of mixed cell death patterns, with cell death in isolated single cells (e.g. **Fig 5A** white outlined blue arrows, 48 hpi,) and in GFP+ ballooning cells which rapidly converted from GFP expressing giant cells to dead cell clusters, indicated by uptake of the red cytotox fluorescent dye (e.g. **Fig 5A** white outlined black arrows, **Fig 5B,C**). Kinetic analysis of some GFP+ ballooning cells revealed characteristics consistent with pyroptosis, including cell rounding and swelling followed by bubble-like membrane protrusions, suggestive of osmotic swelling prior to lysis (**Fig 5B-C, Video S3-5**) [75, 76]. Notably, the profound morphological changes observed with high-dose infection were significantly blunted in Nec-1 treated cultures (right column, **Fig 5A**), which showed increased GFP expression and increased cell death relative to intermediate dose infection, but with a significant paucity of GFP+ large swollen cells when compared to high dose infection alone (**Video S6-7**). The pronounced morphological changes observed with high dose infection were readily observed across multiple fields and infected wells (**Fig S5, Video S3-5**). High dose infection was characterized by the greatest amount of green fluorescence, with Nec-1 treatment characterized by reduced green fluorescence that was still above that observed with intermediate dose infection (**Fig 5D**). In these infected cell cultures, the overall magnitude of cell death, indicated by red fluorescence, was relatively comparable between high dose infection with or without Nec-1 (**Fig 5E**). When quantifying the area of images with high green and high red fluorescence detection, Nec-1 treated cultures showed an increase relative to high dose infection alone beginning by 24 hpi, equalized by 48 hpi (**Fig 5F**). In total, these data demonstrated the time-dependent impact of high dose infection of J774 macrophage cell morphology, a process characterized by induction of large, ballooning GFP+ cells followed by cell death in isolated cells and cell lysis in a fraction of large swollen cells. Among these processes, Nec-1 treatment profoundly blunted the generation of GFP+ ballooning cells, suggesting that Nec-1 blocks morphological changes induced during late stages of high dose infection.

**Figure 5:**
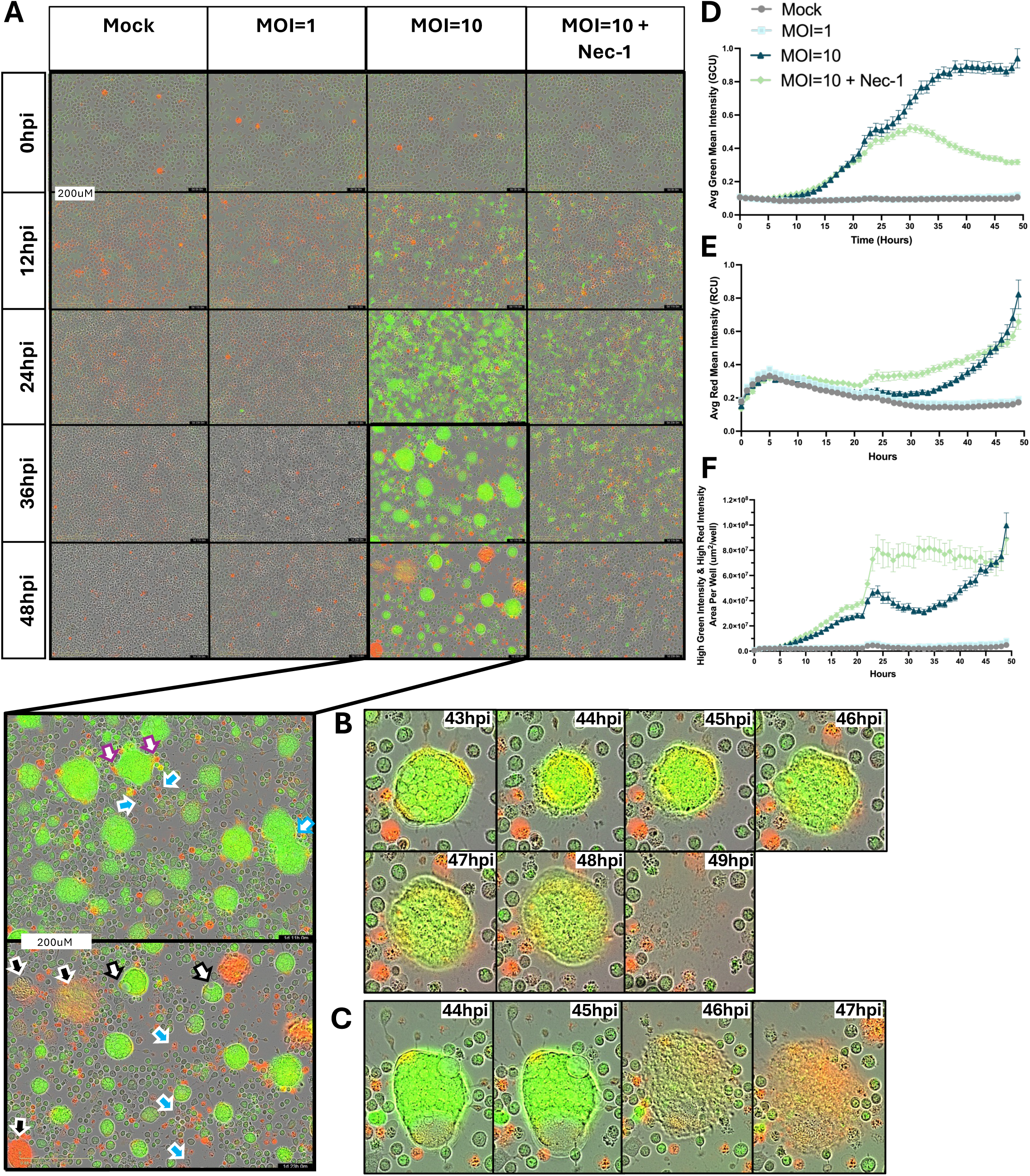
High dose infection of J774 macrophages reveals dramatic changes in cell morphology overtime. Incucyte-based kinetic analysis of MHV68.H.GFP infection in J774 macrophages comparing mock, MOI=1,MOI=10 and MOI=10+Nec-1 treatment over 49 hours. (A) Representative images of J774 macrophages infected with MHV68.H.GFP at indicated MOI with or without 30 µM Nec-1 treatment at 20X magnification. Green fluorescence represents GFP expression by MHV68.H.GFP, with dead cells indicated by red fluorescence (Incucyte Cytotox red dye). Large cells or ballooning cells interacting with or phagocytosing other cells are indicated by purple outlined white arrows. Cells that are merging are indicated by blue outlined white arrows. Large cells with vacuolization are indicated by black outlined white arrows. Cells that have undergone or are undergoing death are indicated by white outlined black arrows. Small cells that are undergoing cell death are indicated by white outlined blue arrows. Images display time in bottom right corner that represents hours since imaging start. Hours post infection (hpi) is indicated on the left column for each row. Enlarged images at bottom left of Figure indicate enlarged views of MOI=10 at 36 (top) and 48 (bottom) hpi. (B-C) Representative high magnification images of large, GFP+ cells undergoing death. Each row follows the fate of an individual large, ballooning cell over the indicated times post-infection. (D) Average green mean intensity (GCU) indicative of GFP expression by the MHV68.H.GFP virus, (E) Average red mean intensity (RCU) indicative of cell death in these cultures, with (F) High green and red intensity (yellow) area per well as defined by non-adherent Incucyte analysis. Data show mean ± SD, with individual symbols showing average measurement of 3 wells from independent biological experiments with 9 images per well. Scale bar indicates 200 microns. Links to Incucyte videos for all conditions provided in Supplemental Video 1-7.

## DISCUSSION

Myeloid cells are an important cell type in yHV infection, representing both a target of infection as well as a key cell type that influences the host response [9, 21, 24]. Despite their importance, the outcomes and mechanisms regulating yHV infection in myeloid cells remains relatively understudied. Here, we sought to investigate the impact of virus infection dose, building on our recent findings where we reported pronounced differences in infection outcome that occur in an MOI-dependent manner using the MHV68 system [20]. While our studies focused on reductionist system to modulate infection dose, it is important to note that the amount of virus that a macrophage is exposed to in vivo will be heavily influenced by its proximity to productively infected cells in primary infection or reactivation.

By comparing high and intermediate infection doses, we demonstrated that high dose infection is associated with significantly decreased cell viability by 48 hpi, coinciding with lytic replication. While lytic replication is associated with cell death, notably, the magnitude of cell death observed with high infection dose greatly exceeded that observed in other infection conditions resulting in virus replication in macrophages (e.g. IL-4 treatment). Specific inactivation of viral DNA [47] and inhibition of the viral polymerase show that high dose infection-associated cell death requires intact viral DNA and virus transcription, which occurred independently of viral DNA replication or late gene transcription. These findings strongly suggest that viral envelope or tegument proteins are not sufficient for high dose infection-associated death, and is more likely due to expression of immediate early and/or early viral genes. Though the viral genes that drive this process remain to be defined, we find that the vBcl2, a known inhibitor of apoptosis and autophagy [34, 77–79] limits cell death in high dose infection; we also find that trehalose-induced autophagy and Nec-1 can profoundly attenuate this cell death.

The yHVs encode a suite of cell death antagonists, including vBcl2 homologs [34, 77–79] and vMAP (a mitochondrial anti-apoptotic protein [80]), with demonstrated crosstalk between apoptosis and autophagy [81]. Previous studies demonstrate complex interaction with caspases [82, 83], but rule out caspase 1/11-mediated GSDMD activated inflammatory cell death [84]. Despite the benefit of Nec-1, a necroptosis inhibitor, biochemical analysis of cell death strongly implicates that high dose infection does not result in necroptosis, but elicits apoptosis, potentially followed by gasdermin E-mediated secondary pyroptosis. While the genetic requirement for GSDME remains to be verified in future studies, it is intriguing to note that there is precedent for this model in other viruses, including herpes simplex virus (HSV)-2 induced apoptosis and GSDME-induced neuronal cell death [42], an observation also recently made in enterovirus 71 infection [85].

Kinetic analysis of high dose infection cultures revealed an unanticipated complexity, with MOI- and time-dependent cell death accompanied by the formation of large, virus-infected, multinucleated ballooning cells, a morphology restricted by Nec-1. GFP+ ballooning cells often appeared to interact with neighboring dying cells. Notably, a fraction of these swollen giant cells progressed to lysis, with morphological features consistent with secondary pyroptosis and dramatic lysis by 48 hpi, that resembles HSV-2 induced GSDME activation [42]. How these giant, ballooning cells form is unknown but could represent cell fusion events (a process reminiscent of virus-induced syncytia [86] or multinucleated giant cells formation [87]), an inflammatory response to pyroptosis in nearby cells [88], a virus driven response to promote replication as spread [89], or phagocytosis and/or efferocytosis (the process of apoptotic body uptake ([90]), particularly as this phenomenon occurs in a macrophage cell line. Whether a comparable process occurs during yHV infection in vivo (e.g. resulting in the generation of virus-laden, giant, swollen, lysis-prone cells) remains an important unanswered question.

During our investigation of potential cell death mechanisms, we used necrostatin-1, a compound first identified as a necroptosis inhibitor [61, 62]. Although Nec-1 reduced cell death and cytotoxicity, further analysis of cell death was not consistent with necroptosis (absence of phospho-MLKL and resistance to RIPK3 inhibition), raising the possibility that Nec-1 may function by inhibiting RIPK1-dependent apoptosis ([65, 91]) or through another mechanism. Notably, longitudinal analysis of Nec-1 treated cultures revealed that while cell death was relatively comparable with high dose infection conditions, Nec-1 treated cultures had pronounced reduction in the formation of large, virally-infected, ballooning cells. These data imply that the protective effects of Nec-1 may be due to an altered response to cell death, rather than regulating cell death itself. While the underlying mechanism(s) of Nec-1 protection in this context remain to be defined, our studies used the original Nec-1 compound, an inhibitor of RIPK1 [61] and IDO [67, 68]. Future studies will investigate the relative contributions of these molecules and the possibility that Nec-1 is regulating either formation of giant cell development or efferocytosis, each with the potential to affect inflammatory cytokine production. While Nec-1 specificity is unclear in this setting, use of Nec-1 and a host of related compounds [92] holds great promise in a range of inflammatory conditions [62, 93, 94]. The potential of these treatments during yHV infection and disease is yet to be tested.

This study has multiple limitations. First, studies rely on reductionist in vitro studies to examine the outcome of MHV68 infection, with many studies based on the J774 cell line. While we demonstrated that high dose infection resulted in cell death in primary peritoneal macrophages, the impact of Nec-1 was solely observed in J774 cells. This could be due to distinct mechanisms of death or the specific microenvironment of high dose infection of J774 cells, in which a monoculture of cells results in the formation of burgeoning, virus-infected cells that expand over time. Macrophages also demonstrate pronounced phenotypic variation, highly influenced by tissue microenvironment; it remains to be determined how conserved the response to high dose infection is across multiple macrophage subsets. Second, inhibitor studies primarily rely on a limited number of concentrations, with at least one of the inhibitors (i.e. Nec-1) known to have more than one target of action. Third, it remains to be determined how these pathways and findings translate to the human yHVs (i.e. EBV, KSHV) and to in vivo infection (in mice or in humans). To address the question of virus dose head-on, future studies will need to develop methods to either modulate or measure virus dose in infected macrophages in vivo.

In total, our studies demonstrate that high dose infection of macrophages by MHV68 induces a pronounced series of alterations characterized by extensive cell death, lytic virus replication, and the development generation of large, virus-infected ballooning cells that can succumb to a terminal lytic burst. These infection outcomes can be regulated by the vBcl2, by the induction of autophagy and by Nec-1, although the precise stages regulated by these treatments remains to be determined. These data reveal the multifaceted interactions between yHV infection and macrophages and identify potential pathways to be therapeutically targeted to alter how macrophages respond to yHV infection, with the potential to alter the host inflammatory response.

## METHODS

### Viruses, Cells and Tissue Culture

Wild-type (WT) MHV68, MHV68.LANA::βlac [48, 95], MHV68.H.GFP [40], MHV68.ORF59.GFP [96], MHV68.C-RTA [46], MHV68.vBcl2.stop [69], MHV68.vBcl2.AAA [35], and MHV68.vCyc.stop [97] viruses were grown and prepared as described previously [40, 48, 95], with virus titers determined from at least three independent plaque assays.

Mouse 3T12 fibroblasts (ATCC CCL-164) and J774A.1 macrophages (“J774”; ATCC TIB-67), and Mouse Embryonic Fibroblasts (MEFs, generated as in [98] were cultured in complete DMEM (cDMEM): Dulbecco’s modified Eagle medium (DMEM; Life Technologies, Cat No. 11965-092) supplemented with 5% (3T12) or 10% fetal bovine serum (J774, MEFs) (FBS; Atlanta Biologicals), 2 mM L-glutamine, 10 U/mL penicillin, and 10 μg/mL streptomycin sulfate (Gibco, Cat No. 10378016) at 37°C with 5% CO_2_.

J774 macrophages were plated at 4-5×10^5^ cells/mL in 6- or 12-well plates. 3T12 fibroblasts were plates at 2×10^5^ cells/mL in 12-well plates. MEFs were plated at 1.5×10^4^ cells/well in 96 well plates supplemented with Gibco™ Amphotericin B (Thermo Fisher Scientific, Cat. No. 15290018). Peritoneal cells were harvested by peritoneal lavage, following intraperitoneal injection of 10 mL of ice cold non-additive DMEM into eight-week old female C57BL/6J mice (stock #000664) obtained from the Jackson Laboratory (Bar Harbor, ME) and were maintained in a specific-pathogen-free environment.

### Drug Treatments

Cells were treated with various drug treatments, with timing of administration illustrated in **Fig S6**. Cell death inhibitors: Media was replaced with 10% cDMEM with Necrostatin-1 (30 µM [99]; SelleckChem, Cat No. S8037) or LDC7559 (1 µM [63]; SelleckChem, Cat No. S9622) 2h prior to infection, with UH15-38 (500 nM, 750 nM, or 1 µM [66]; Tocris, Cat No. 8104) 1h prior to infection, or MCC950 (500nM, Invivogen Cat No. inh-mcc) 30 min prior to infection, replenished with fresh media and inhibitor at 1hpi. Antibody blocking TNFα (25 µg/mL) (clone XT3.11-CP054; BioXcell, Cat No. BE0058) or Rat isotype control (25 µg/mL) (BioXcell, Cat No. BE0088) was added 2h prior to infection, with antibody concentrations maintained in media during and after infection.

Cell death inducers: To induce pyroptosis, cells were resuspended and plated in LCIS (Thermofisher, Cat No. A59688DJ) + LPS (1 µg/mL; from *E. coli* O55:B5, Sigma-Aldrich, Cat No. L2880) for 4h, followed by addition of Nigericin (20 µM: Sigma-Aldrich, Cat No. N7143) for an additional 1h [100]. To induce necroptosis, cells were resuspended and plated in LCIS + z-VAD-FMK (50 µM; SelleckChem, Cat No. S7023) + sm-164 (5 µM; SelleckChem, Cat No. S7089) [101] for 1h, followed by addition of recombinant mouse TNFα (25 ng/mL; ThermoFisher, Cat No. 315-01A) for an additional 4h before harvest [102]. To induce autophagy, cells were treated with Trehalose (100mM; Sigma, Cat No. T9531) 2h prior to infection [34]. To inhibit virus replication, cells were treated with cidofovir [(*S*)-1-(3-hydroxy-2-phosphonylmethoxypropyl) cytosine; SelleckChem, Cat No. S1516], added 1 hpi at the indicated concentration, replenished at 24hpi. To induce lytic infection in macrophages, cells were pre-treated with recombinant mouse interleukin-4 (IL-4; 10ng/mL [20]; PeproTech, Cat. No. 214-14) 16h prior to infection. Cells were collected for flow cytometry, LDH or plaque assay at 48 hpi.

### Centanamycin-mediated Virus Inactivation

To inactivate MHV68 viral DNA, WT MHV68 (or other reporter viruses) was incubated with 100 µM centanamycin (CM, kindly provided by Dr. Moses Lee [47]) at room temperature for 2 hours, after which CM-treated virus was diluted in a final volume of 25 mL DMEM, followed by virus concentration by 39,800xg centrifugation for 2 hours at 4° C (Beckman Coulter, Avanti J-25, JA-17 rotor). Virus pellets were resuspended in the original volume and stored at -80° C. Vehicle control samples were processed identically in the absence of CM and titered by plaque assay to define any loss of virus titer due to incubation and centrifugation steps (as in **Fig S2A**). CM treatment of MHV68 at 4° C failed to fully inactivate MHV68 at either 1 or 100 µM (**Fig S2D**).

### Infection

MHV68 infections were based on live cell counts with the TC20 Automated Cell Counter (Bio-Rad) with Trypan Blue dye (Bio-Rad, Cat. No. 145-0021) to calculate viral inocula, with multiplicity of infection (MOI) defined as PFU/cell. Cells were incubated at 37°C in 5% CO2 for 1 hour, with rocking every 15 min, before removal of viral inoculum and replacement with 1mL of cDMEM in 12-wells. Samples were harvested at the indicated hours post-infection (hpi). For MEF infections, virus was serially diluted to 1 PFU, 0.33 PFU, 0.11 PFU or 0.037 PFU per well, with each dilution measured in 24 wells, monitored for CPE over 10 days.

### Plaque Assay

Plaque assay quantification of viral titer was performed using 3T12 cells, as in [20]. Briefly, cells were plated in 12-well plates at 8.5×10^4^ cells per well one day prior to infection. At time of infection, media was removed from cells, infected with virus containing samples for 1 h at 37°C at 5% CO2, with plates rocked every 15 min prior to overlay with a 1:1 mix of 10% cDMEM and carboxymethyl cellulose (CMC; Sigma, Cat. No. C-4888) supplemented with Gibco™ Amphotericin B (Thermo Fisher Scientific, Cat. No. 15290018). Plates were incubated for 8 days before staining with 0.5% methylene blue to count plaques. Samples were tittered using 10-fold dilution series, diluted in 5% cDMEM; an internal plaque standard of known titer was included in each plaque assay to ensure reproducible sensitivity.

### Light Microscopy

At the indicated time points, 12-well plates of J774 cells were imaged on an AMG EVOS FL (ThermoFisher) using a 20X phase objective to capture brightfield or green fluorescence images. Immediately following image acquisition, samples were harvested and processed for flow cytometric or LDH analysis. To image nuclei, cells were fixed with 4% PFA (paraformaldehyde, ThermoFisher, Cat No. J6899.AK) and stained with DAPI (200 nM; 4’,6-Diamidino-2-Phenylindole, Dilactate, Biolegend, Cat No. 422801) following manufacturer’s protocol, visualized using a 20X phase objective on EVOS FL.

### Flow Cytometry

J774, 3T12 cells, or primary peritoneal cells were harvested at the indicated times, were stained with LIVE/DEAD fixable Near-IR Dead Cell (Invitrogen, Cat. No. L10119) at a 1:1500 dilution in BSS wash (111mM Dextrose, 2mM KH2PO4, 10mM Na2HPO4, 25.8 CaCl2•2H2O, 2.7mM KCl, 137mM NaCl, 19.7mM MgCl•6H_2_O, 16.6mM MgSO_4_). To assess LANA.βlac expression in MHV68 LANA::βlac infected samples, cells were then incubated with CCF2-AM (Thermo Fisher, Cat. No. K1023) at a 1:10 dilution following the manufacturer’s protocol, quantifying the frequency of cells with beta-lactamase mediated substrate cleavage. To analyze cell survival in primary peritoneal macrophages infected ex vivo, peritoneal cells were stained with fluorescently conjugated antibodies against F4/80 (1:400; clone BM8, ThermoFisher, Cat No. MF48021) and CD11b (1:400; clone M1/70, BioLegend, Cat No. 101220) in the presence of Fc receptor-blocking antibody (Clone 2.4G2, Cytek, Cat No. 70-0161-U100), and fixed in 1% paraformaldehyde prior to collection on an Agilent Novocyte Penteon flow cytometer. All flow cytometry experiments included unstained, single-stain, and full-minus-one controls to define background fluorescence, fluorescent signal spread, and compensation.

### Lactate Dehydrogenase (LDH) Measurement

J774 macrophages were plated at 5×10^5^ cells/mL in 12-well plates in 10% cDMEM. 1 hour before indicated harvest time points, cells were scraped, counted and diluted in LCIS to 2.5×10^4^ cells/mL, with 50 µL transferred to a 96-well plate. Quantitation of lactate dehydrogenase (LDH) release was measured by absorbance using Invitrogen CyQUANT LDH Cytotoxicity Assays kit (ThermoFisher, Cat No. C20300) according to the manufacturer’s instructions. % Cytotoxicity (LDH release) was determined from spontaneous and maximum LDH absorbance measurements according to manufacturer’s instructions.

### Incucyte Live-Cell Imaging Analysis

For Incucyte analysis, J774 cells were infected in a 12-well plate at indicated MOI for 1 hour before viral inoculum was removed and replaced with 1mL of cDMEM with 250nM Incucyte® Cytotox Red Dye (Sartorius, Cat No. 4632), followed by imaging by Sartorius Incucyte S3 Imaging system (Sartorius, Cat No. 4647). Cells were imaged every 1hr at 20X magnification. Using Incucyte Analysis software (Incucyte 2023A GUI) we applied the Non-Adherent Cell analysis setting to quantify green (a measure of MHV68.H.GFP expression) or red fluorescence (Cytotox Red) calibrated units (GCU or RCU respectively) over time. To quantify total yellow intensity per well, cell-by-cell analysis was applied to the non-adherent cell set. Data was plotted using the average values from 9 separate fields of view per well, from separate three wells from each of three independent biological replicates, with error bars indicating standard deviation defined by Incucyte software.

### Western Blot

J774 cells subjected to various treatments were collected and centrifuged at 3000xg for 3 min. Samples were resuspended in 100 µL Triton lysis buffer (1% SDS, 1 mM EDTA, 0.01 M Tris pH 8, and 150 mM NaCl), PMSF (Roche, catalog #10837091001) and calyculin A (Sigma Aldrich, catalog #208851) as in [73], incubated on ice for 5 min prior to centrifugation at 3000xg for 3 min for collection of the Triton soluble fraction. Samples were then boiled at 95° C for 5 min in Laemmli sample buffer (BioRad, Cat No. 1610747).

Samples were prepped as in [73]. Lysates and supernatant were separated using SDS-PAGE, and proteins were wet transferred onto 0.22 μm nitrocellulose membranes. Membranes were then washed with Tris-buffered saline with 0.01% Tween-20 (TBST) and incubated with primary antibody diluted in 5% bovine serum albumin in TBST for 16 hours at 4°C or 1-2 hours at 25°C. Membranes were washed thrice in TBST for 5 minutes then incubated with HRP-conjugated secondary antibody diluted in 5% milk in TBST for 1-2 hours at 25°C. Membranes were again washed thrice in TBST then developed with enhanced chemiluminescence.

Primary antibodies were used at 1:1000 dilution, with clones and commercial sources in **Table S1**. Secondary antibodies were used at 1:10,000-1:5000 dilution: anti-rabbit (Cell Signaling, Cat No. 7074), anti-mouse (Cell Signaling, Cat No. 7076).

### Statistical Analysis and Software

Data analysis, graphing and statistical analyses were done in GraphPad Prism (version 11.0.2; GraphPad Software, San Diego, California USA, www.graphpad.com).

Flow cytometry data were analyzed using FlowJo (version 10.10.1. Ashland, OR: Becton, Dickinson and Company; 2026). Statistical significance was tested by unpaired, nonparametric, Mann-Whitney t test or 1-way ANOVA, detailed in each figure legend.

## Ethics Statement

All animal studies were performed in accordance with the recommendations in the Guide for the Care and Use of Laboratory Animals of the National Institutes of Health. Studies were conducted in accordance with the University of Colorado Denver/Anschutz Institutional Animal Use and Care (IACUC) Committee, in compliance with Animal Welfare Assurance of Compliance policy (assurance no. D16-00171), with studies approved under IACUC protocol #189. All procedures were performed under isoflurane anesthesia, and all efforts were made to minimize suffering.

## ACKNOWLEDGEMENTS

We thank Dr. Anh Le of the Cell Technology Shared Resource and Dr. Joshua Jackson of Sartorius for consultation and assistance with Incucyte analysis. We thank Christine Childs and Kristina Terrell of the University of Colorado Cancer Center Flow Cytometry Shared Resource for training and assistance with flow cytometry. For generously sharing recombinant viruses, we thank Dr. Craig Forrest (MHV68.HygroGFP), Dr. Craig Forrest and Dr. Laurie Krug (MHV68.ORF59.GFP), and Dr. Ren Sun (MHV68.C-RTA). This study was supported by the National Institutes of Health (NIH) grant R01 AI157201 awarded to LvD and ETC, R35 GM141603 awarded to DAW, Molecular Biology T32 pre-doctoral training grant (NIH5T32GM136444) awarded to REK, NIH grants F31 AI176728 and T32 AI052066 awarded to EAS, with additional support by the National Institutes of Health Cancer Center Support Grant (P30CA046934), including the Cell Technologies Shared Resource (RRID: SCR_021982) and the Flow Cytometry Shared Resource (RRID: SCR_022035) of the University of Colorado Cancer Center.

## AUTHOR CONTRIBUTIONS

Conceptualization: G.V., D.W.A., L.F.v.D., E.T.C.;

Methodology: M.L., D.W.A.;

Validation: G.V., G.N.R., R.E.K, E.A.S., D.W.A.;

Formal Analysis: G.V., E.T.C.;

Investigation: G.V., G.N.R., R.E.K, E.A.S., D.W.A.;

Data Curation: G.V., E.T.C.;

Writing – original draft: G.V., L.F.v.D., E.T.C.;

Writing – review & editing: G.V., L.F.v.D., E.T.C.;

Visualization: G.V., L.F.v.D., E.T.C.;

Supervision: L.F.v.D., E.T.C.;

Project administration: G.V., L.F.v.D., E.T.C.;

Funding acquisition: L.F.v.D., E.T.C.

## DECLARATION OF GENERATIVE AI AND AI-ASSISTED TECHNOLOGIES IN THE WRITING PROCESS

Generative AI was not used in preparation of this manuscript.

---

**Supplementary Table 1.** Antibodies for Western Blotting.

| Antibody | Commercial Source | Catalog # and RRID |
| --- | --- | --- |
| Rabbit polyclonal anti-Caspase-3 | Cell Signaling Technology | Cat# 9662; RRID:AB_331439 |
| Rabbit monoclonal anti-Caspase-8 | Cell Signaling Technology | Cat# 4790; RRID:AB_10545768 |
| Rabbit polyclonal anti-Caspase-8 (cleaved) | Cell Signaling Technology | Cat# 9429; RRID:AB_2068300 |
| Mouse monoclonal anti-Caspase-9 | Cell Signaling Technology | Cat# 9508; RRID: AB_2068620 |
| Rabbit monoclonal anti-GSDMD (mouse) | Abcam | Cat# ab209845; RRID: AB_2783550 |
| Rabbit monoclonal anti-GSDME | Abcam | Cat# ab215191; RRID: AB_2737000 |
| Rabbit monoclonal Antibody anti-MLKL | Cell Signaling Technology | Cat #37705; RRID: AB_2799118 |
| Rabbit monoclonal Antibody anti-PARP | Cell Signaling Technology | Cat# 9532; RRID: AB_659884 |

## SUPPLEMENTAL FIGURE LEGENDS

**Supplemental Figure 1:**
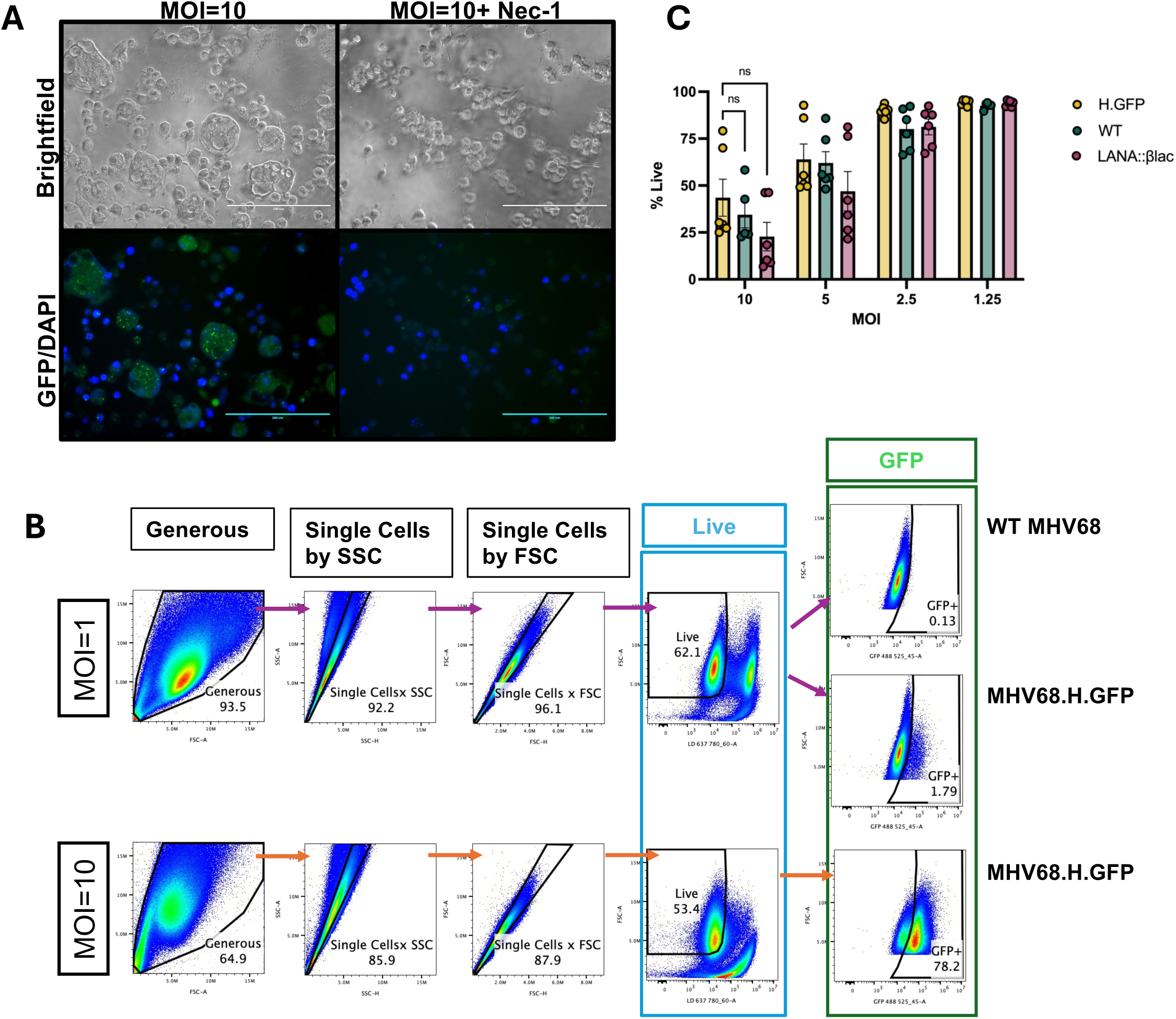
Virus induced cell death of J774 macrophages is highly dose-dependent. (A) 20X brightfield and corresponding GFP/DAPI images of J774 macrophages infected with H.GFP virus at an MOI = 10 and analyzed for GFP expression with DAPI (blue) at 48 hpi. Scale bar indicates 200 µm. (B) In support of Figures 1-4. Flow cytometry-based gating strategy to define the frequency of cells that were live and/or GFP positive, analyzing MHV68.H.GFP-infected J774 macrophages at a MOI = 1 or 10. Cells were analyzed by flow cytometry, using sequential focused analysis (”gating”) to quantify the frequency of single cells (defined as ”Single Cells by SSC” or “Single Cells by FSC”) that are viable (“Live”, boxed in blue, fourth column from the left), defined by exclusion of a dye that identifies dead cells. Live cells were then analyzed for GFP fluorescence (”GFP”, boxed in green, right column), with background defined based on fluorescence detected in the GFP channel following infection with WT MHV68 (top plot, right column). Controls included unstained and samples singularly positive for fluorescence using the LIVE/DEAD NearIR dye or cells infected with MHV68.H.GFP without additional fluorophores. (C) In support of Figure 1. J774 macrophages were infected with MHV68.H.GFP, WT MHV68 or MHV68.LANA::βlac virus at indicated MOI (PFU/cell). Frequency of live cells was determined by flow cytometry at 48 hpi. Data show mean ± SEM, with individual symbols showing independent biological samples from 3 independent experiments with 2 biological replicates per experiment. Statistical analysis was done using Mann-Whitney test with statistically significant differences indicated, ns (not significant). Flow cytometry gating strategy for primary peritoneal cells and for MHV68.LANA::βlac-infected samples shown in Supplemental Figure 2B, 3B.

**Supplemental Figure 2:**
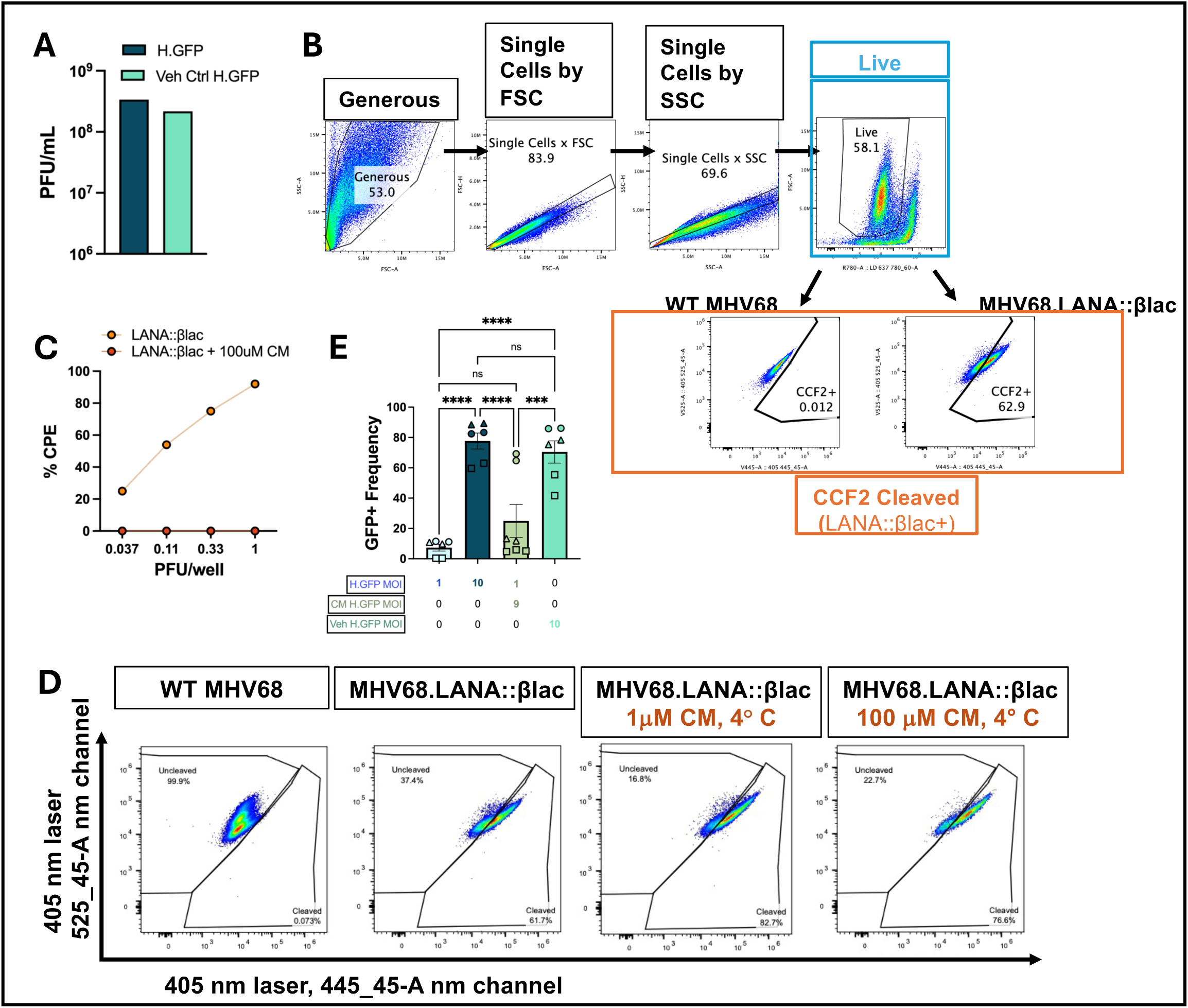
Effect of centanamycin on MHV68 infection and gene expression. In support of Figure 2. (A) Virus titer defined by plaque assay, comparing virus stock treated with vehicle control (Veh Ctrl H.GFP), to quantify the effect of sample processing during centanamycin (CM) incubation on virus concentration, compared with unmanipulated virus stock (H.GFP). (B) In support of Fig 2B-C. Gating strategy for live and LANA::βlac+ cells in MHV68.LANA::βlac infected 3T12 fibroblasts at an MOI = 1. Cells were analyzed by sequential focused analysis (”gating”) to quantify the frequency of single cells (defined as “Single Cells by FSC” or” Single Cells by SSC”) that were viable (Live boxed in “blue”) followed quantitation of the frequency of cells with evidence of cleaved beta-lactamase substrate (measured by fluorescence excited by the 405 nm laser, with emission in the 445 nm/45 nm detection filter (“CCF2 Cleaved” in orange). Controls included WT MHV68 infected samples incubated with CCF2 substrate, to define background fluorescence, as well as single stain controls for CCF2 substrate cleavage or live/dead stain (LIVE/DEAD NearIR). (C) Virus infectivity of CM-treated virus, defined by quantifying the percentage of wells demonstrating cytopathic effect (CPE) on highly sensitive mouse embryonic fibroblasts (MEFs), comparing dilution series of untreated MHV68.LANA::βlac (i.e. LANA::βlac) with CM-treated virus stock (i.e. LANA::βlac + 100 µM CM at 22° C) at 10 dpi (24 replicate wells/virus/condition). This condition (i.e. 2 hour incubation of virus with 100 µM CM at 22° C) was used for all virus inactivation studies in Fig 2A-F. (D) Quantification of viral gene expression as a function of CM treatment, assessing the frequency of cells with CCF2 substrate cleavage among single, live cells, a measure of LANA.βlac expression, by flow cytometric analysis of 3T12 cells at 48 hpi (MOI=1 PFU/cell). Data depict samples incubated with CM at 4° C, conditions that failed to inactivate MHV68. Images depict representative flow cytometry-based analyses, assessing CCF2 substrate cleavage (defined by cells with positive fluorescent signal in the 405 nm, 445_45 nm detection channel) among single, live cells from 3 independent experiment with 2 biological replicates per experiment. (E) Frequency of live J774 cells infected with H.GFP with or without CM treatment, harvested at 48 hpi, defined by flow cytometry, with shapes indicating paired experimental samples. Data show mean ± SEM, with individual symbols showing independent biological samples from 3 independent experiment with 2 biological replicates per experiment. Statistical analysis was done using two-way ANOVA (ordinary, with Tukey’s multiple comparisons test), with statistically significant differences indicated, ***p<0.001,****p<0.0001, ns (not significant).

**Supplemental Figure 3:**
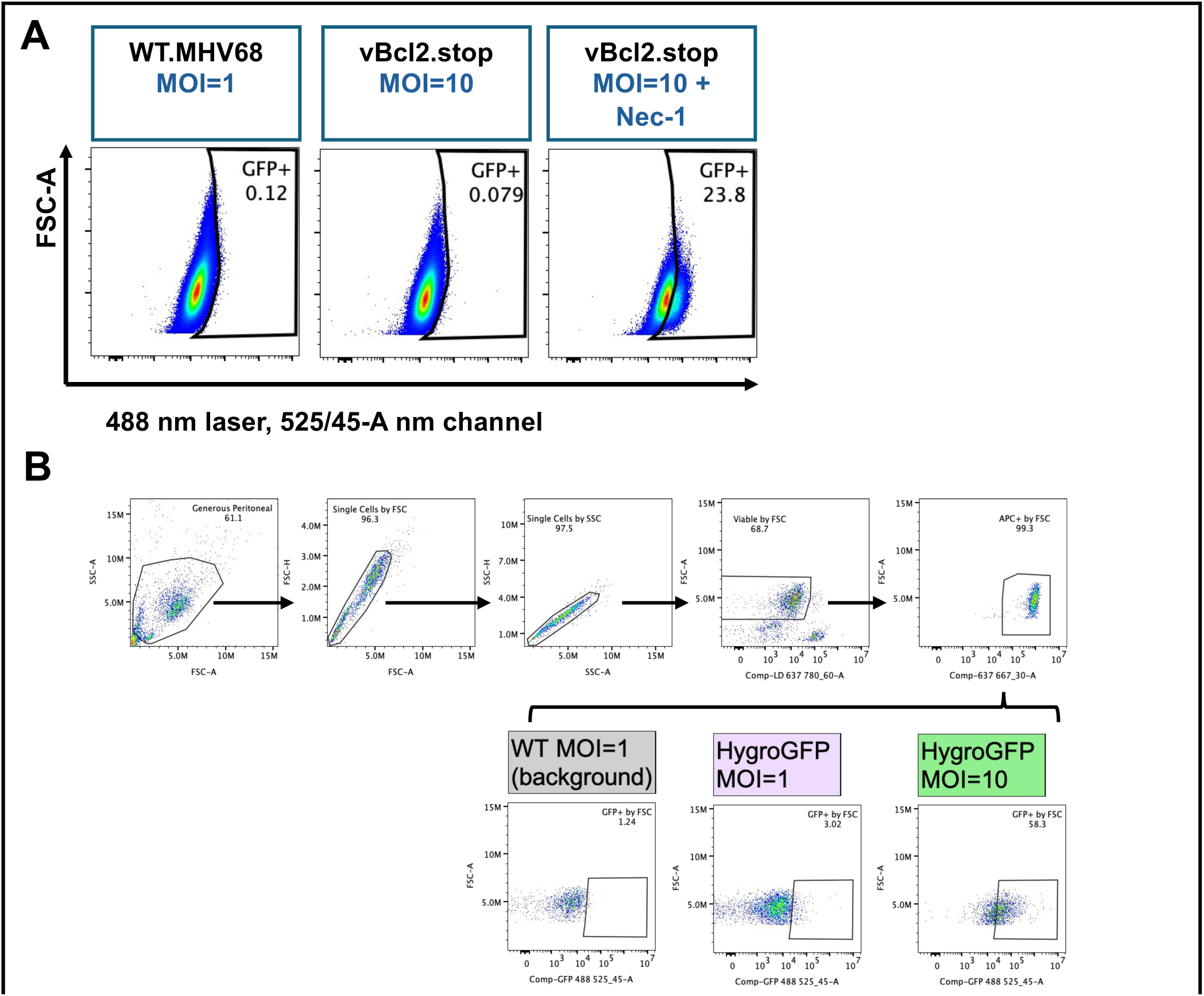
Necrostatin-1 treatment increases autofluorescence in MHV68-infected J774 cells. (A) In support of Figure 3D, 3G. J774 macrophages were infected with WT MHV68 or MHV68.vBcl2.stop at an MOI = 1 or 10 with or without Nec-1 treatment (30 µM). Fluorescence was analyzed by flow cytometry at 48 hpi, with data showing cells that were viable, single cells. Data from 1 independent experiment with 2 biological replicates per condition. (B) In support of Fig 3G-H. Gating strategy to assess viability and GFP expression in primary peritoneal macrophage following ex vivo infection. Data show gating hierarchy to define the frequency of viable, single, CD11b+ F4/80+ cells, and the frequency of GFP+ viable, single, CD11b+ F4/80+ cells. Bottom row depicts representative flow cytometry plots for WT MOI=1 (to define background fluorescence), compared to MOI=1 or MOI=10 infection with HygroGFP virus.

**Supplemental Figure 4:**
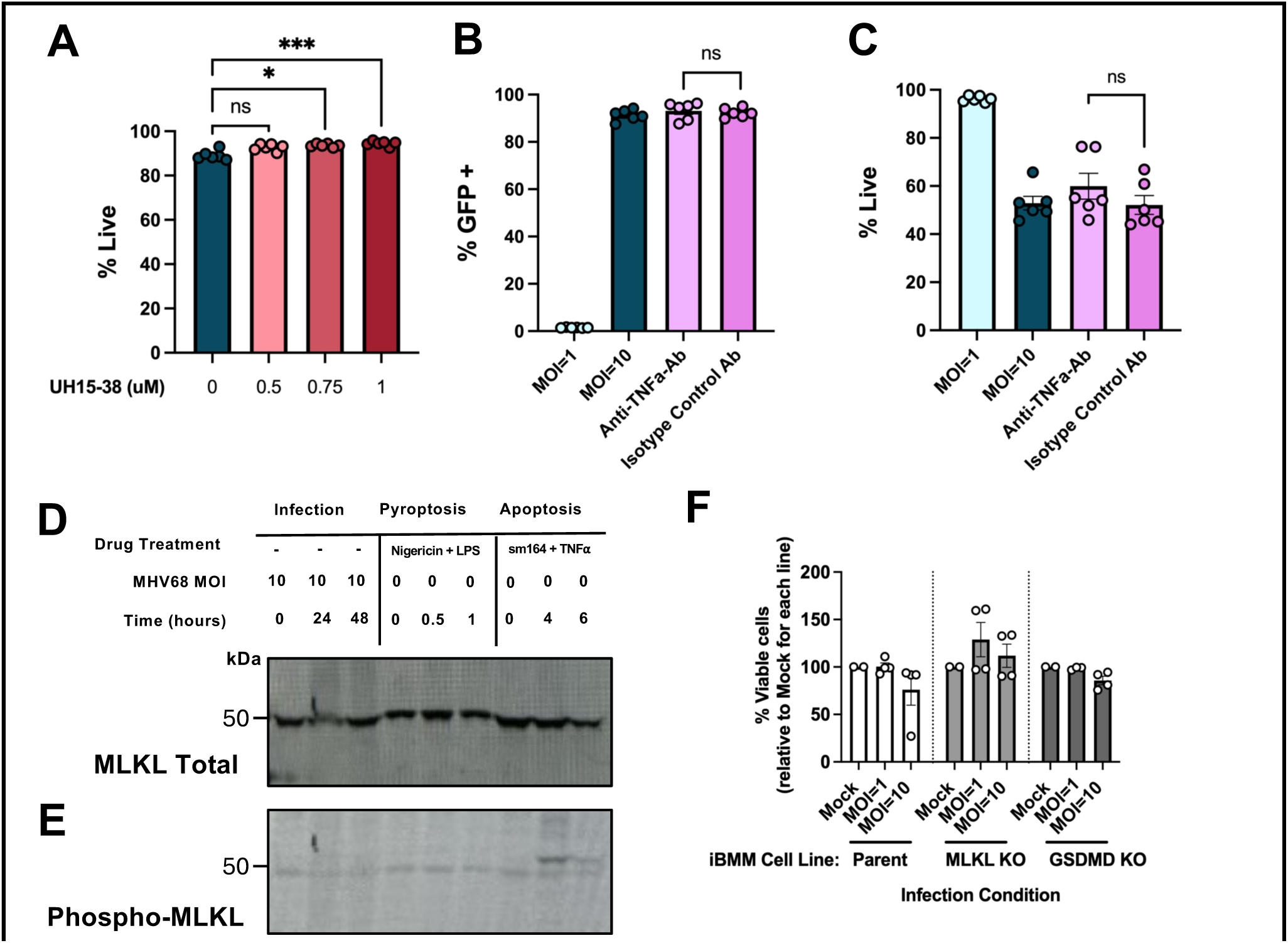
Impact of different interventions on J774 cell viability and the outcome of high dose infection. In support of Figure 3-4. (A) J774 macrophages were treated with UH15-38 at indicated concentration 1h prior to ‘mock’ infection with 10% cDMEM + UH15-38 for 1h followed by maintenance of UH15-38 in media. Percent live cells was measured by flow cytometry 48 hours post-“mock infection”. (B-C) J774 macrophages were treated with no drug, Anti-TNFα-IgG or Anti-Rat-IgG (25 µg/mL) 2h prior to infection with MHV68.H.GFP. Treatment was maintained through infection to harvest at 48 hpi in which (B) the frequency of GFP expressing cells or (C) the frequency of viable cells were defined by flow cytometry. (D) Immunoblot analysis of cell lysates from infected J774 cells (left) or J774 cells stimulated with pyroptosis inducers (nigericin with LPS, middle) or apoptosis inducers (sm164 with TNFα, right), probed with an antibody to MLKL in support of Fig 4G, or (E) probed with an antibody to phospho-MLKL; blot images shown are representative of three independent experiments with consistent results. (F) Analysis of cell survival in MHV68-infected immortalized bone marrow macrophages (iBMMs), comparing mock infection with MHV68 infection using an MOI=1 (1 PFU/cell) or MOI=10. Studies analyzed cell viability (defined by flow cytometry as the frequency of viable, single cells), in parental, MLKL-deficient (MLKL KO), or GSDMD-deficient (GSDMD KO) iBMMs. Viability was defined relative to mock infected cultures for each cell line for each experiment. Data from two to three independent biological experiments, with each experiment done in one to two parallel culture wells plated from a single culture, infected, processed and analyzed independently. Data show mean ± SEM, with individual symbols depicting data from all culture wells. Statistical analysis was done using Mann-Whitney test (B, C) comparing isotype versus blocking antibody or one-way ANOVA analysis (A, Kruskal-Wallis test subject to Dunn’s multiple comparisons test) compared the effect of UH15-38 dose relative to no drug treatment. Statistically significant differences indicated, *p<0.05, ***p<0.001, ns (not significant).

**Supplemental Figure 5:**
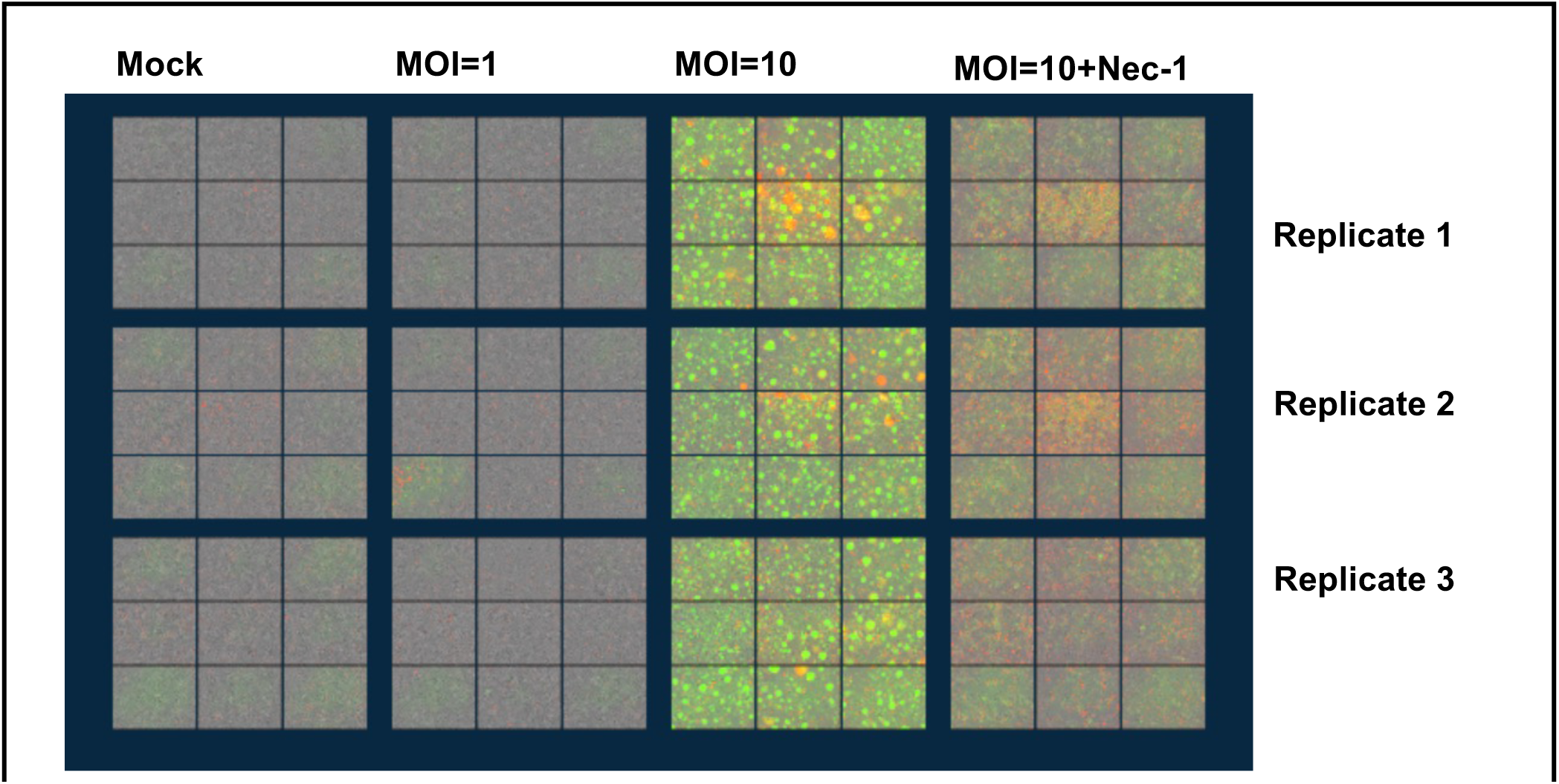
Microscopy images of MHV68 high dose infection of J774 cells show distinct fluorescent features. In support of Figure 5. Overview of Incucyte collected images of J774 cells subjected to the four indicated conditions (mock, MOI=1, MOI=10, MOI=10+Nec-1) analyzed at 48 hpi. Each condition had 3 replicates (1 per row) with data collected from 3 replicates per condition and 9 fields of view per replicate per condition per timepoint.

**Supplemental Figure 6:**
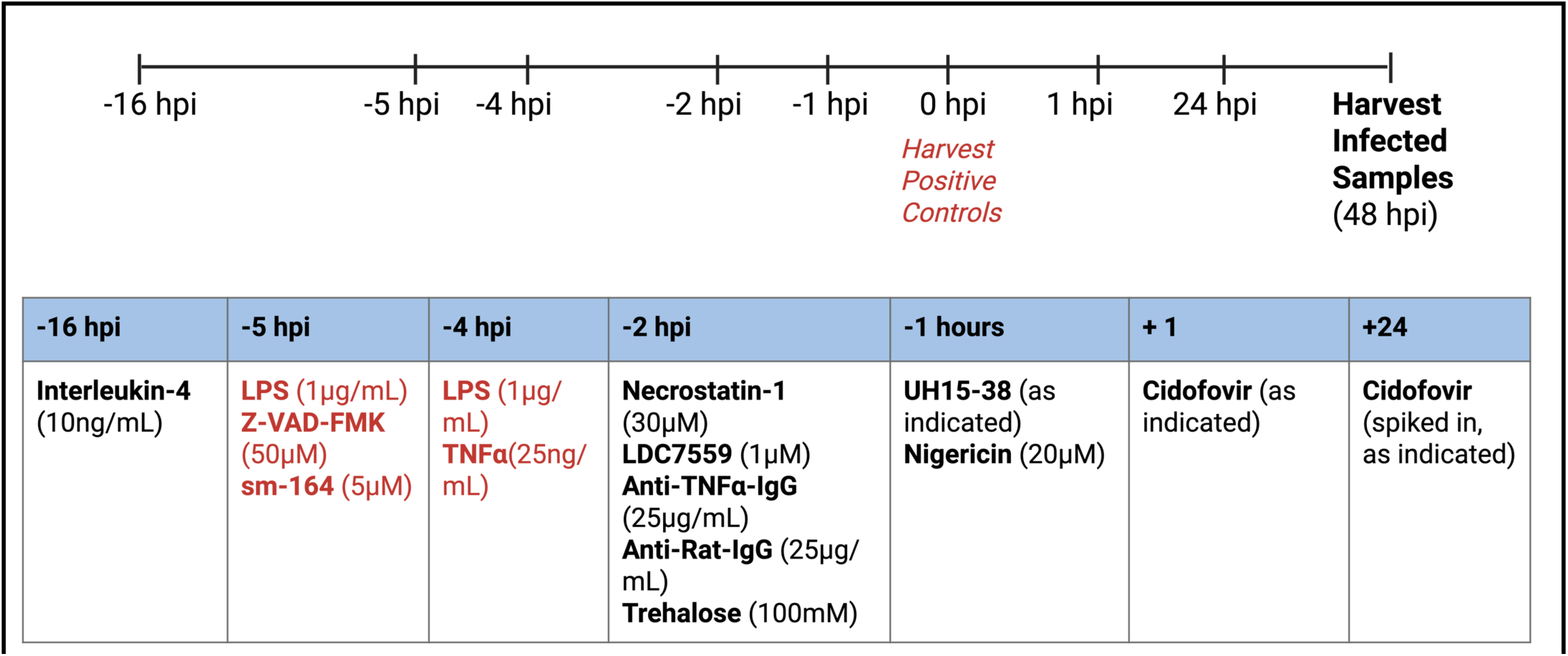
Treatment and dosing schedule of compounds used to study the outcome of MHV68 infection in macrophages. hpi, hours post-infection.

Supplemental Videos 1-7 (Videos S1-S7): Kinetic analysis of infection dynamics in MHV68-infected J774 cells. In support of Fig 5. Video time course over 1-49 hpi, J774 cells treated as indicated, with Cytox Red cell death indicated by red fluorescence, virus gene expression indicated by GFP green fluorescence, scale bar measure = 200 microns. * in file name denotes videos containing large GFP+ cells swelling and ballooning before dramatic lysis visible between 19-25 seconds.

| <u>Supp Video #</u> | <u>File name</u> | <u>Infection condition</u> | <u>Drug</u> | <u>Video link</u> |
| --- | --- | --- | --- | --- |
| Video S1 | Mock_A1_6 | none | none | Mock_A16.mp4 |
| Video S2 | MOI=1_A2_2 | MOI=1 | none | MOI1_A22.mp4 |
| Video S3 | MOI=10_B3_3 * | MOI=10 | none | MOI10_Rep1_B33.mp4 |
| Video S4 | MOI=10_A3_1 * | MOI=10 | none | MOI10_Rep2_A31.mp4 |
| Video S5 | MOI=10_C3_9 * | MOI=10 | none | MOI10_Rep3_C39.mp4 |
| Video S6 | Nec-<br>1_MOI=10_C4_4 | MOI=10 | Nec-1 | MOI10_Nec1_Rep1_C44.mp4 |
| Video S7 | Nec-<br>1_MOI=10_B4_2 | MOI=10 | Nec-1 | MOI10_Nec1_Rep2_B42.mp4 |

Supplemental Video 1 (Video S1): J774 cells mock treated. Video time course shows 1-49 hpi. Cell death events are indicated by Cytox red color. Scale bar measures 200 microns. Filename: Mock_A1_6. Culture: Uninfected J774 cells. <u>Mock_A16.mp4</u>

Supplemental Video 2 (Video S2): J774 cells infected with MHV68.H.GFP at a MOI of 1. Video time course shows 1-49 hpi. Cell death events are indicated by Cytox red color. Virus gene expression is indicated by GFP green color. Scale bar measures 200 microns. Filename: MOI=1_A2_2. Culture: J774 cells infected at intermediate MOI=1 dose. <u>MOI1_A22.mp4</u>

Supplemental Video 3 (Video S3): J774 cells infected with MHV68.H.GFP at a MOI of 10. Video time course shows 1-49 hpi. Cell death events are indicated by Cytox red color. Virus gene expression is indicated by GFP green color. Scale bar measures 200 microns. Filename: MOI=10_B3_3. Culture: J774 cells infected at high MOI=10 dose, dramatic swollen/ballooning GFP+ cells and lysis between 19-25 seconds. <u>MOI10_Rep1_B33.mp4</u>

Supplemental Video 4 (Video S4): J774 cells infected with MHV68.H.GFP at a MOI of 10. Video time course shows 1-49 hpi. Cell death events are indicated by Cytox red color. Virus gene expression is indicated by GFP green color. Scale bar measures 200uM. Filename: MOI=10_A3_1. Culture: J774 cells infected at high MOI=10 dose, dramatic swollen/ballooning GFP+ cells and lysis between 19-25 seconds. <u>MOI10_Rep2_A31.mp4</u>

Supplemental Video 5 (Video S5): J774 cells infected with MHV68.H.GFP at a MOI of 10. Video time course shows 1-49 hpi. Cell death events are indicated by Cytox red color. Virus gene expression is indicated by GFP green color. Scale bar measures 200 microns. Filename: MOI=10_C3_9. Culture: J774 cells infected at high MOI=10 dose, dramatic swollen/ballooning GFP+ cells and lysis between 19-25 seconds. <u>MOI10_Rep3_C39.mp4</u>

Supplemental Video 6 (Video S6): J774 cells infected with MHV68.H.GFP at a MOI of 10 + 30 µM necrostatin-1 treatment. Video time course shows 1-49 hpi. Cell death events are indicated by Cytox red color. Virus gene expression is indicated by GFP green color. Scale bar measures 200 microns. Filename: Nec-1_MOI=10_C4_4. Culture: J774 cells infected at high MOI=10 dose in presence of Necrostatin-1. MOI10_Nec1_Rep1_C44.mp4

Supplemental Video 7 (Video S7): J774 cells infected with MHV68.H.GFP at a MOI of 10 + 30 µM necrostatin-1 treatment. Video time course shows 1-49 hpi. Cell death events are indicated by Cytox red color. Virus gene expression is indicated by GFP green color. Scale bar measures 200uM. Filename: Nec-1_MOI=10_B4_2. Culture: J774 cells infected at high MOI=10 dose in presence of Necrostatin-1. MOI10_Nec1_Rep2_B42.mp4

